# Exosomal Profiling Reveals Mechanisms of Hibernation-Associated Neuroprotection

**DOI:** 10.64898/2026.06.23.733742

**Authors:** Francisco M. Nadal-Nicolás, Rachel McNeel, Kirsten Overdahl, Alan Jarmusch, Kiyoharu J. Miyagishima

## Abstract

Glaucoma is a group of eye diseases that affects 4 million people in the US and is one of the leading causes of vision loss due to damage to the eye’s optic nerve (ON) which is composed of axons from retinal ganglion cells (RGCs) that transmit visual information to the brain. Injury to the ON often triggers RGC death and subsequent loss of visual function. Despite its increasing prevalence worldwide, effective therapies for glaucoma remain elusive. Notably, the thirteen-lined ground squirrel (TLGS) exhibits intrinsic neuroprotection during hibernation; however, reproducing this protective state pharmacologically has proven challenging. To elucidate the metabolic mechanisms underlying this resilience, we conducted untargeted metabolomic analyses on TLGS retinas at 6 hours, 3 days, and 7 days following ON crush. Retinas from awake and hibernating animals were compared to identify temporal and state-dependent metabolic signatures. Distinct metabolomic profiles were observed in hibernating animals relative to their awake counterparts. Pathway analyses revealed coordinated regulation of amino acid, lipid, and purine metabolism that likely contributes to hibernation-induced resilience. Furthermore, our findings indicate that hibernating TLGS retinas increase exosome biogenesis, prompting *in vitro* validation using TLGS-derived exosomes, which demonstrated robust neuroprotective and anti-inflammatory effects. Proteomic and transcriptomic characterization of exosomal cargo identified conserved miRNAs, mRNAs, and proteins implicated in redox balance, cytoskeletal stabilization, and stress-response regulation. Collectively, these data support the hypothesis that metabolic reprogramming and exosome-mediated intercellular signaling underlie hibernation-associated neuroprotection. Modulating these pathways may provide a blueprint for novel therapeutic strategies to mitigate neurodegeneration and promote recovery following optic nerve injury.

**Graphical Abstract:** Illustration depicting state-dependent metabolic responses to optic nerve crush (ONC) injury in Thirteen-lined Ground Squirrels (TLGS). In Awake animals, injury triggers enhanced ATP production through the TCA cycle, leading to excessive reactive oxygen species (ROS) generation and subsequent retinal ganglion cell (RGC) death. In contrast, Hibernating animals shift toward lipid metabolism and utilize ATP for the biosynthesis of ceramides and sphingolipids, promoting membrane integrity and exosomal signaling. Additionally, a range of metabolites associated with hibernation-linked neuroprotection are elevated, contributing to enhanced RGC survival.

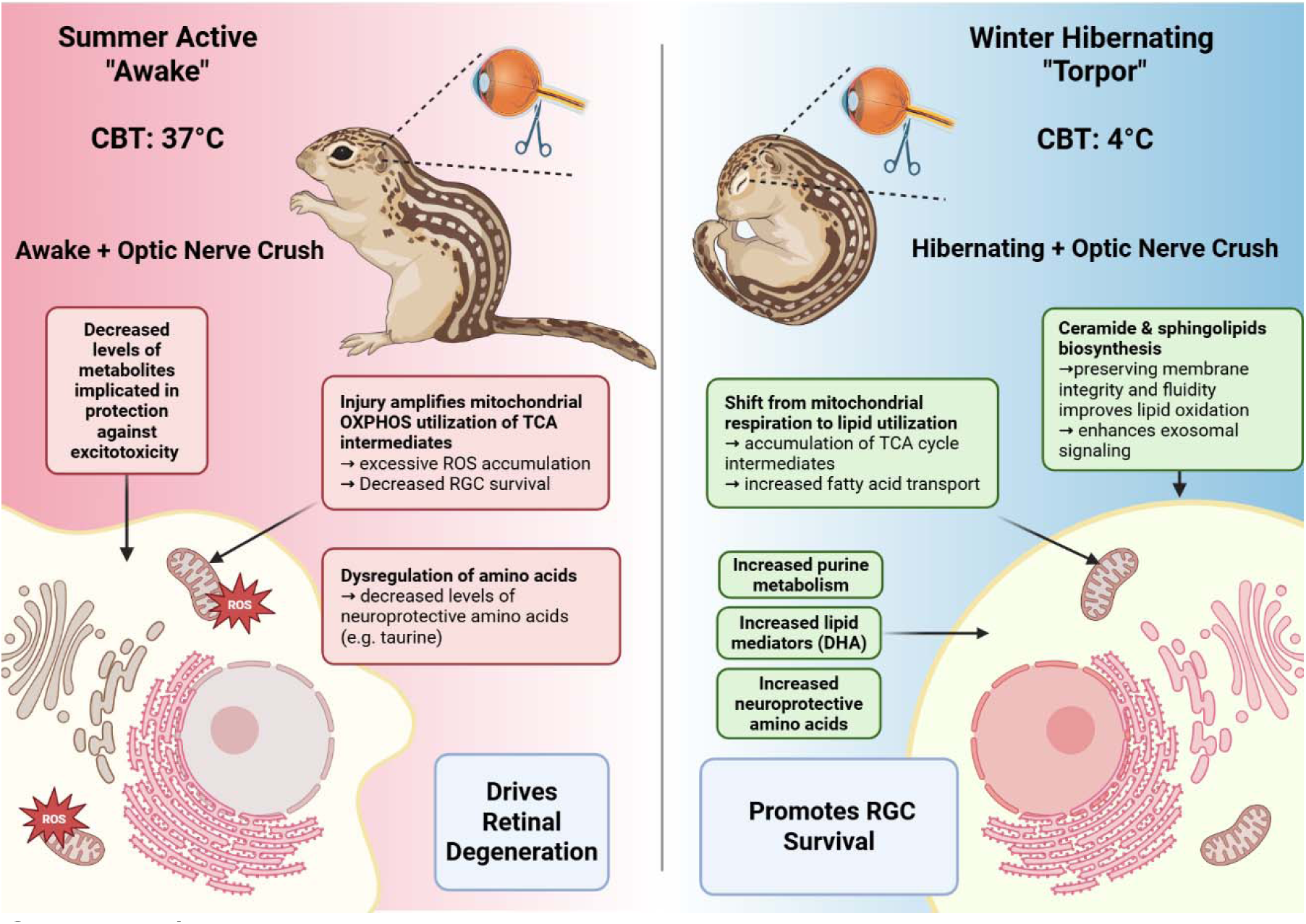

## Introduction

The precise pathophysiology of glaucoma remains incompletely understood, hampering the development of effective therapies. Several mechanisms have been proposed, including mechanical compression injury resulting from elevated intraocular pressure, ischemic damage to the retinal ganglion cell (RGC) axons, impaired ocular blood flow, genetic susceptibility, autoimmune-mediated axonal degeneration, and neurodegenerative processes that may originate in the brain. Together, these factors contribute to both primary and secondary neuronal degeneration. Given the devastating and irreversible nature of vision loss due to glaucoma, there is a critical need for innovative strategies to prevent RGC death, preserve visual function, and promote neural repair and recovery.

One promising line of investigation arises from the study of mammalian hibernators, such as the thirteen-lined ground squirrel (TLGS), *Ictidomys tridecemlineatus*, which has evolved exceptional mechanisms of cellular preservation. During torpor, these animals profoundly suppress metabolic activity, reduce oxidative stress, and attenuate inflammation [1, 2]. Remarkably, they endure prolonged periods of hypothermia and ischemia - conditions that mimic aspects of glaucoma pathogenesis, including hypoxia, energy depletion, and oxidative stress – without sustaining irreversible neuronal damage. During deep hibernation, core body temperature (CBT) can fall from 37°C to as low as 4°C, yet they avoid the cellular damage typically associated with such conditions, making hibernators a powerful model for studying endogenous neuroprotective mechanisms.

To explore this phenomenon, we employed a controlled ON crush (ONC) model to simulate axonal damage due to glaucoma [3, 4]. In awake TLGS, this injury induces axonal degeneration and progressive RGC loss. However, RGCs in hibernating ground squirrels exhibit intrinsic resistance to injury, suggesting an evolutionarily conserved neuroprotective mechanism that may inform new therapeutic strategies. Indeed, hibernating ground squirrels naturally suppress these pathological cascades during torpor, thereby preserving RGC viability despite ON injury [5–11]. To better understand the mechanisms that confer genuine neuroprotection in hibernation, we performed untargeted metabolomic profiling of retinas from TLGS subjected to ONC injury while either awake or in hibernation. Retinal samples were collected at 6 hours (acute phase post-injury), 3 days (peak microglial activation), and 7 days (onset of homeostatic transition, with approximately 50% RGC loss [7, 12]).

This temporal and physiological comparison revealed that hibernation is associated with increased exosome biogenesis and secretion—metabolic signatures consistent with enhanced extracellular vesicle production. Building on these findings, we demonstrate that exosomes derived from hibernating retinas exert potent neuroprotective and anti-inflammatory effects, reducing neuronal death induced by cold stress and suppressing lipopolysaccharide (LPS)-evoked inflammatory responses. Together, these results identify exosome-mediated communication as a key component of hibernation-associated neuroprotection and a promising therapeutic avenue for the treatment of traumatic optic neuropathies.

## Results

### Neuroprotection by Hibernation and Metabolic Inhibition After Optic Nerve Crush

The physiological state of hibernation (**Figure 1A-C**) profoundly influences the retinal response to optic nerve crush (ONC) injury (**Figure 1D**). Consistent with our previous findings [8–11], we observed that retinal ganglion cells (RGCs) in the nasal retina remain largely intact in hibernating ground squirrels after partial ONC. In contrast, awake animals exhibited substantial RGC degeneration in the same region highlighting the intrinsic neuroprotective advantage conferred by the hibernation state.

**Figure 1.**
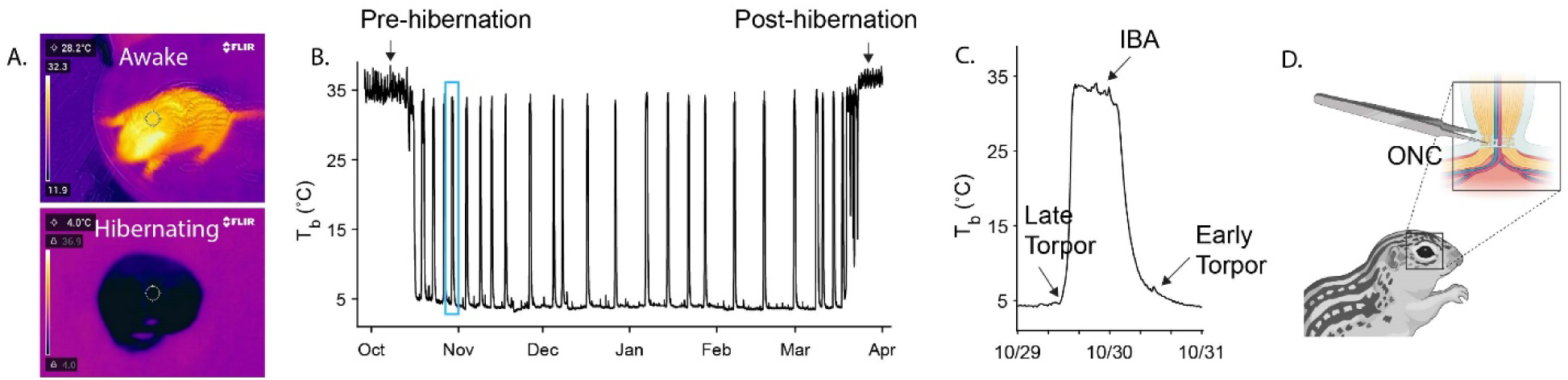
Hibernation protects retinal ganglion cells (RGCs) after optic nerve crush (ONC) in thirteen-lined ground squirrels (TLGS). (A) Thermal images of an awake (top) and hibernating (bottom) TLGS illustrate the marked drop in surface body temperature during torpor during which time the animals are kept in a hibernaculum at 4°C to mimic the low winter temperatures of their native Wisconsin. (B) The body temperature (T_b_) of a TLGS plotted over the course of a year capturing the seasonal changes in temperature that occur during the hibernation season. The blue box highlights an interbout arousal during which the animal awakens from torpor and recovers body temperature to 37°C periodically for approximately 24-48 hours. (C) Zoomed in image of an interbout arousal (IBA) showing the spontaneous rapid recovery to 37°C and the controlled decrease in body temperature to 4°C during the entrance into torpor. (D) Schematic of the ONC procedure in TLGS.

### Temporal Dynamics of Microglial and Astrocytic Responses at the Optic Nerve Lesion Site

The progression of RGC degeneration is accompanied by pathological changes in the optic nerve (ON) at the crush site, ultimately contributing to visual function loss [13]. **Figure 2A** shows representative confocal images of the crush site from ON sections of Awake and Hibernating animals at time points corresponding to those used for retinal analysis (**Figure S4**). Iba1, a marker for resident microglia, is upregulated upon activation, noticeably increased at 3 days post injury (ONC 3d) and further at 7 days (ONC 7d) in Awake animals. This activation pattern is mirrored by expression of CD68, a lysosomal marker for activated microglia, which is low in naïve (uninjured) animals but elevated at ONC 3d and ONC 7d in the same groups.

**Figure 2.**
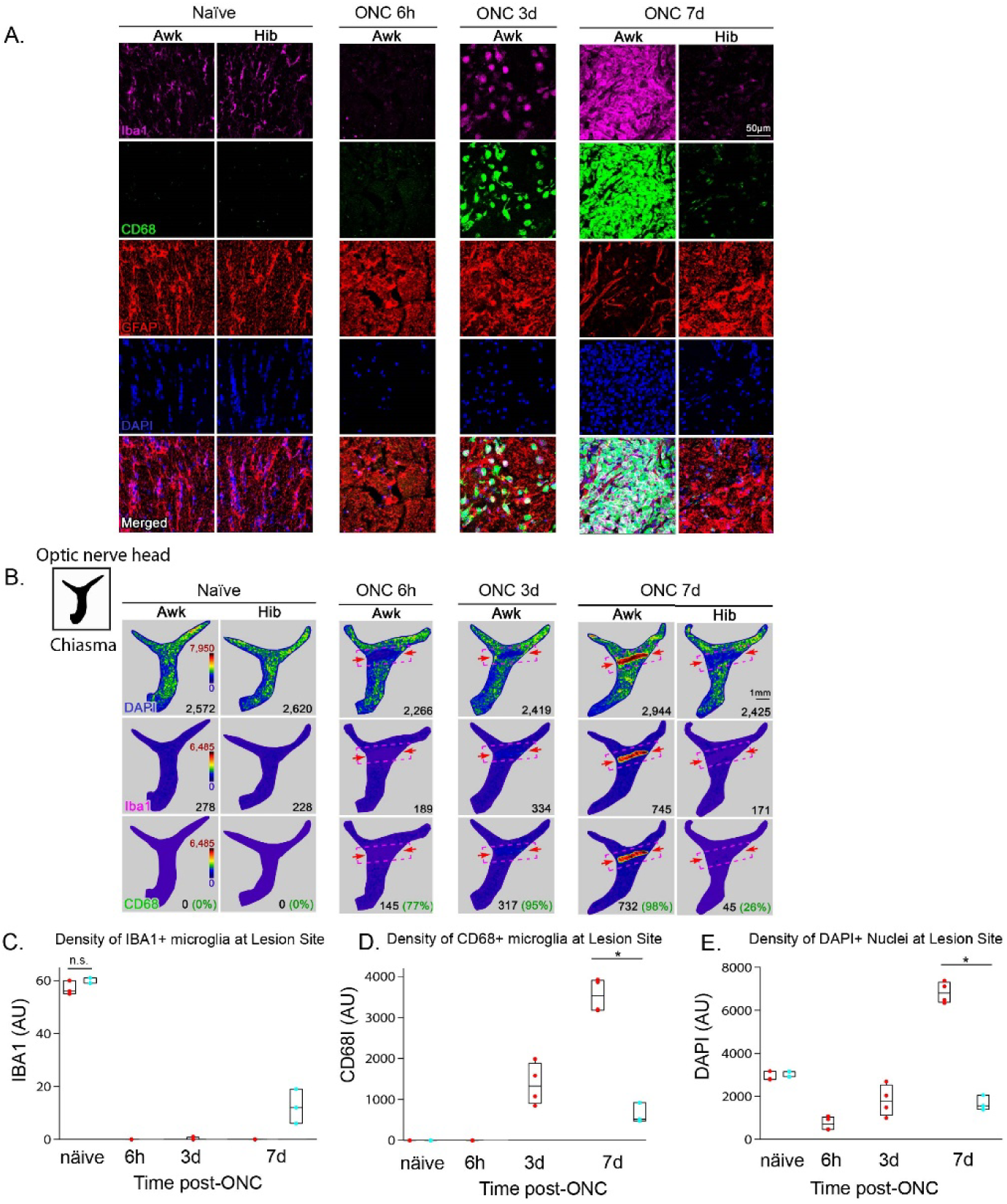
Images and analysis of optic nerves corresponding to the retinas used for untargeted metabolomic profiling. (A) Representative high-magnification images of the optic nerve crush (ONC) site in Awake and Hibernating animals under naïve conditions and at multiple time points post-injury (6 hours, 3 days, and 7 days). Optic nerves are immunolabeled for Iba1 (microglia, magenta), CD68 (activated microglia, green), and DAPI (nuclei, blue). Immunostaining reveals dynamic remodeling of the glial environment during the progression of injury and repair. Merged images reveal that the Hibernating condition at 7 days post-ONC closely resembles the naïve state. Scale bar: 50 µm. (B) Representative isodensity plots showing the spatial distribution of DAPI, Iba1, and CD68 immunolabeling across the different time points post-injury. Each isodensity plot represents the density of the immunofluorescence marker indicated in the bottom left corner of the first image on the left. The top row shows DAPI (nuclei) density, the middle row shows Iba1 (microglia) density, and the bottom row shows CD68 (activated microglia) density. The region of the crush site, indicated by red arrows, is delineated by a red dashed box superimposed on the isodensity plot. The inset in the upper left of Figure 2B shows the orientation of the horizontal optic nerve indicating the optic nerve head, the crush site, and the axons extending to the chiasma. Scale bar: 1mm. (C) Quantification of IBA1/CD68 microglia at the lesion site reveals a time-dependent decrease following ONC. (D) In contrast, CD68 microglia show a progressive increase over time post-injury. Notably, optic nerves from hibernating squirrels maintain low levels of CD68 cells even at 7 days post-ONC. (E) DAPI nuclei count initially decline immediately after injury but subsequently increase as cells infiltrate the lesion site. This cellular infiltration is absent in the optic nerves of hibernating animals at 7 days post-ONC. n.s. indicates no significant difference, while significant changes are denoted by asterisks (*P ≤ 0.05)

Notably, Iba1 and CD68 expression in Hibernating animals at ONC 7d closely resembles that of the naïve condition, indicating reduced microglial activation. In parallel, glial fibrillary acidic protein (GFAP, red), a marker of astrocytes, is rapidly upregulated within the first 6 hours post-injury in Awake animals – reflecting an early astrogliotic response. However, GFAP expression subsequently declines at ONC 3d and ONC 7d, consistent with astrocyte degeneration near the lesion. By ONC 7d, astrocytes are notably absent from the crush site, although they are expected to repopulate this area at later stages of gliotic remodeling [14]. In contrast, GFAP expression in Hibernating animals at the lesion site mirrors the early post-injury increase seen in Awake animals (ONC 6h).

DAPI staining, used to label nuclei, shows increased intensity at the lesion site by ONC-7d compared to naïve animals, suggestive of cellular infiltration or proliferation. These DAPI+ nuclei co-localize with Iba1 (magenta), confirming their identity as microglia. Isodensity plots of entire optic nerves (**Figure 2B**) clearly demarcate the crush site at ONC 3d and ONC 7d in Awake animals, with dense accumulations of CD68+ and Iba1+ microglia indicating activation. Quantification revealed a time-dependent decline in cells expressing only Iba1 (**Figure 2C**), suggesting a shift toward a more activated phenotype co-expressing CD68. Accordingly, integrated density measurements confirmed increased CD68 expression at the lesion site post-injury (**Figure 2D**). In contrast, CD68 levels in Hibernating animals remained low even at ONC 7d, aligning with the reduced accumulation of microglia and corresponding DAPI+ nuclei at the lesion site (**Figure 2E**).

### Comprehensive Metabolomic Profiling Reveals Injury-Associated Metabolic Shifts

Using untargeted metabolomic analysis, we identified 218 metabolites in negative ionization mode and 230 metabolites in positive ionization mode. Together, these datasets provide comprehensive coverage of the retinal metabolite profile and reveal dynamic changes associated with ON injury in the retinas of hibernating and awake animals.

Partial Least Squares Discriminant Analysis (PLS-DA) was used to assess group-specific retinal metabolic profiles across different treatment conditions. The PLS-DA plots represent discriminate analyses between the uninjured and injured TLGS within two dimensions and each point in the PLS-DA plot represents an individual TLGS retina. The mass spectrometry (MS) data is collected in both positive and negative ionization modes. In the negative ionization mode, minimal separation was observed between uninjured and injured awake TLGS across time points indicating similar metabolic characteristics among these groups (**Figure 3A**). In contrast, in the positive ionization mode, uninjured control samples exhibited distinct separation from the injured TLGS groups, indicating injury-associated metabolic differences between the control and the injured TLGS tissues (**Figure 3B**). The hibernating control samples were distinctly separated from the injured hibernating animals in both (negative and positive) ionization modes, suggesting that injury disrupts the metabolome in a manner not accounted for by hibernation alone (**Figure 3C-D**).

**Figure 3.**
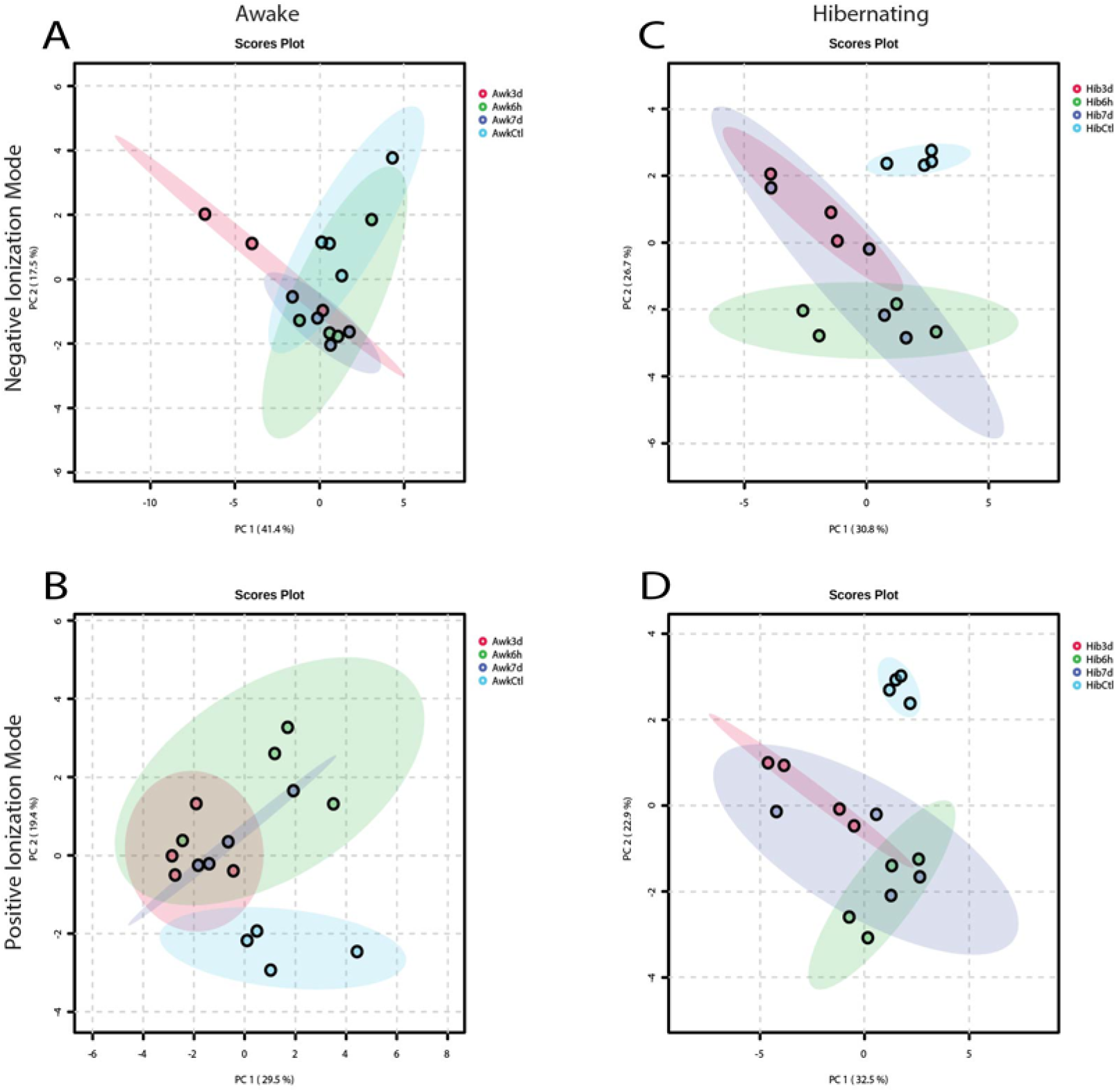
Partial Squares Discriminant Analyses (PLS-DA) plots were used to identify clusters of retinal metabolites in intact and injured awake TLGS and hibernating TLGS under negative ionization mode (A, C) and positive ionization mode (B, D).

### Negative Ionization Mode Metabolomics Reveals Neuroprotective Metabolic Adaptations

MS data from the negative ionization mode were first analyzed by grouping the TLGS retina samples (biological replicates) according to the following experimental conditions: awake and hibernating. Hierarchal clustering analysis of the retinal metabolites in uninjured and injured TLGS revealed metabolites with significant changes in abundance (|log(Fold Change)|>1.5), identifying potential biomarkers of ON degeneration and candidate metabolites for neuroprotection [15].

One cluster, denoted as Group A (yellow), exhibited a significant number of altered amino acids across experimental groups (**Figure 4A**). In Awake squirrels, metabolites in this cluster began at elevated levels but declined following injury and remained suppressed. In contrast, the same metabolites in the Hibernating group also started at elevated levels, became slightly reduced immediately after injury, but returned to elevated levels at 7 days post-injury. Amino acids levels, in general, are known to fluctuate following ON injury due to disruptions in neurotransmitter release, impaired neuronal function, and oxidative stress associated with demands for more energy [16, 17]. The identification of key Krebs cycle intermediates such as isocitritic acid, citric acid, and malic acid - among significantly changing metabolites in Awake animals further supports the notion of altered energy production following injury. The increased utilization of these intermediates likely contributes to increased ATP production to meet the increased demand in response to injury. The excess ATP released from dying RGCs into the extracellular space has been implicated in activating retinal microglia, which includes Müller cells, leading to secondary injury driving further RGC death [18].

**Figure 4.**
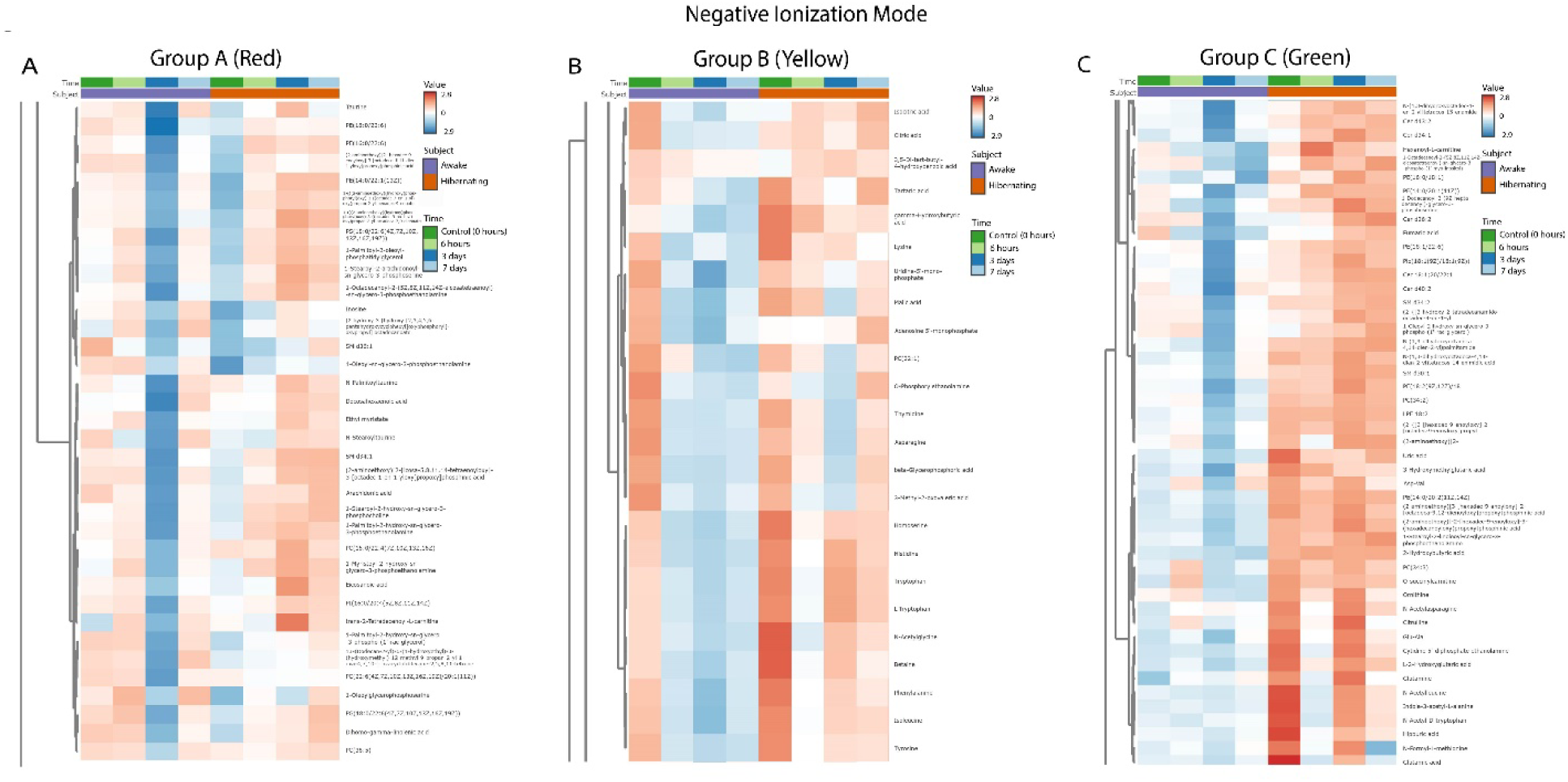
Heatmap clustering analysis of metabolites in the negative ionization mode for the three annotated groups (Group A-red, Group B-yellow, Group C-green). Normalized feature intensity values ranged from -2.9 to 2.8. Subject 1 (S1): awake intact and injured TLGS; subject 2 (S2): hibernated intact and injured TLGS. Time 0 (T0): uninjured, intact control; time 1 (T1): 6 hours post-ONC; time 2 (T2): 3 days post-ONC; time 3 (T3): 7 days post-ONC.

In contrast, in hibernating animals there is an increase in synthesis of vital amino acids such as phenylalanine, tyrosine, and tryptophan. These amino acids are energetically expensive requiring high amounts of ATP to synthesize, and serve as precursors to neurotransmitters like serotonin, which has been found to suppress calcium signaling in retinal axons and reduce glutamate release in the thalamus [19]. Notably, serotonin levels rise in the brains of ground squirrels during hibernation [20]. This utilization of ATP may help to preserve ATP levels and avoid excess ATP-induced injury mechanisms observed in non-hibernating animals (e.g. mice) under similar stress conditions.

Interestingly, histidine, a regulator of HIF-1α [21], remains elevated in hibernating animals after injury but declines in Awake counterparts. Histidine may suppress HIF-1α-induced microglial activation, thereby limiting inflammation. Together, these findings suggest that hibernating squirrels deploy unique metabolic adaptations that prioritize neuroprotection, directing energy toward biosynthetic and anti-inflammatory processes, potentially contributing to their enhanced resilience to optic nerve injury.

The next cluster of metabolites, designated as Group B (green), revealed significant changes in five ceramide species (Cer 18:1;20/22:1, Cer d34:1, Cer d34:2, Cer d36:2, Cer d40:2) and two sphingomyelins (SM d30:1, SM d34:2) across the experimental TLGS groups (**Figure 4B**). This cluster is defined by low metabolite abundance in Awake squirrels, before and after injury, but consistently elevated levels in Hibernating squirrels at all time points. This pattern suggests that these lipids may contribute to pro-survival mechanisms unique to the hibernation state. Ceramides, depending on chain length, can exert distinct effects on cell types essential for brain and eye health [22, 23]. While excessive ceramide accumulation is typically associated with toxicity, metabolic dysfunction, and inflammation—features implicated in neurodegenerative diseases and retinal diseases such as Alzheimer’s, Parkinson’s, and age-related macular degeneration (AMD) [22, 24–26] - a distinct pattern emerges under hibernating conditions [27, 28]. In hibernating squirrels, elevated ceramide and sphingomyelin levels may support a protective, pro-survival phenotype. These lipids play vital roles in maintaining plasma membrane fluidity and mediating cell signaling. Moreover, sphingolipids are integral to the formation of extracellular vesicles such as exosomes, which are upregulated in response to cellular stress – including that encountered during torpor [29, 30]. Exosomes transport biologically active cargo—including proteins, mRNA, and microRNAs—which can mediate neuroprotective and regenerative responses. These findings underscore the context-dependent roles of ceramides and sphingolipids, which may be neurotoxic or neuroprotective depending on physiological state and cellular context.

In addition to sphingolipid differences, we observed significantly elevated levels of glutamine and glutamic acid levels, along with their precursor’s ornithine and citrulline in Hibernating squirrels compared to Awake animals. These metabolites are key components of the glutamine-glutamate cycle, essential for neurotransmitter recycling and RGC function [31, 32]. Ornithine and citrulline, as part of the urea cycle, facilitate the elimination of ammonia, mitigating neurotoxic nitrogen buildup. Other notable metabolites elevated under hibernation include N-acetylleucine, which modulates neuronal excitability and exhibits potential neuroprotection, and lysophosphatidylethanolamine (LPE) 18:2, which protects against glutamate-induced toxicity [33]. Antioxidants such as fumaric acid and uric acid are also increased in hibernating animals; both are known to activate the Nrf2 pathway, scavenge free radicals, and inhibit lipid peroxidation during neuroinflammation. Together, these findings reveal a coordinated metabolic program in hibernating squirrels that integrates lipid signaling, nitrogen detoxification, neurotransmitter homeostasis, and antioxidant defense—suggesting a systems-level adaptation that promotes neuronal survival and enhances resilience to optic nerve injury.

A defining feature of the third cluster, designated Group C (red), is the pronounced decrease in metabolite levels observed at 3 days post-injury in Awake squirrels. In contrast, these same metabolites are elevated in Hibernating squirrels, indicating a distinct metabolic shift during the period corresponding to peak inflammation (**Figure 4C**). Among those increased at 3 days post-ONC are several well-characterized retina metabolites including taurine and arachidonic acid, as well as lesser-known neuroprotective compounds such as inosine and various sphingomyelins (SMs) with varying acyl chain lengths (**Figure 4C**). Prior studies have identified phospholipids, particularly sphingolipids such as SM, as potential biomarkers of RGC degeneration [26, 34]. The inclusion of taurine within this cluster is especially noteworthy, given its increasingly recognized role in protecting against retinal neurodegeneration [35]. Taurine depletion has been shown to cause both photoreceptor and RGC loss, as well as impair the phagocytotic ability of the retinal pigment epithelium (RPE) [36–38]. Therefore, the robust increase in taurine levels observed in Hibernating TLGS retinas at 3 days post-ONC suggests a potential role for taurine in shielding retinal cells from oxidative damage and suppressing microglia-mediated secondary injury [39].

Negative ionization mode metabolomics revealed injury-induced alterations in energy metabolism, inflammation, and neurotransmitter balance, with hibernating squirrels exhibiting protective adaptations absent in Awake animals. Hierarchical clustering identified three major metabolite groups with distinct injury responses across experimental conditions. Group A (yellow) highlighted amino acids and TCA intermediates involved in neurotransmitter cycling and energy production, which declined in Awake animals post-injury but recovered in Hibernators—suggesting preserved metabolic flexibility. Group B (green) comprised sphingolipids and ceramides that were consistently elevated in Hibernators and linked to membrane integrity, vesicle signaling, and neuroprotection, contrasting their known pro-inflammatory roles in disease. Group C (red) revealed a striking increase in neuroprotective metabolites, including taurine, inosine, and sphingomyelins, in Hibernating animals at 3 days post-injury—corresponding with a suppression of peak inflammation. These findings suggest that hibernation preserves critical pathways supporting antioxidant defense, neurotransmitter homeostasis and the capacity to reduce excitotoxicity, and cellular resilience, offering promising targets for therapeutic intervention in optic nerve injury.

### Positive Ionization Mode Reveals Hibernation Metabolic Programs Engage Lipid Mediators, Antioxidants, and NAD+ Biosynthesis to Promote Neuroprotection

Metabolomic analysis of the positive ionization mode was performed by comparing TLGS retina samples from Awake and Hibernating groups. Hierarchical clustering of the resulting heatmap revealed several distinct metabolite clusters based on shared expression patterns.

One cluster resembled Group A (yellow) from the negative ionization mode, characterized by metabolites that declined post-injury in Awake animals but remained stable or elevated in Hibernators (**Figure 5A**). This cluster included amino acids such as isoleucine, tyrosine, tryptophan, and lysine—mirroring trends observed in the negative ionization dataset (**Figure 4A**). Additional metabolites that remained elevated in the Hibernating group included proline, creatine, and norepinephrine (**Figure 5A**).

**Figure 5.**
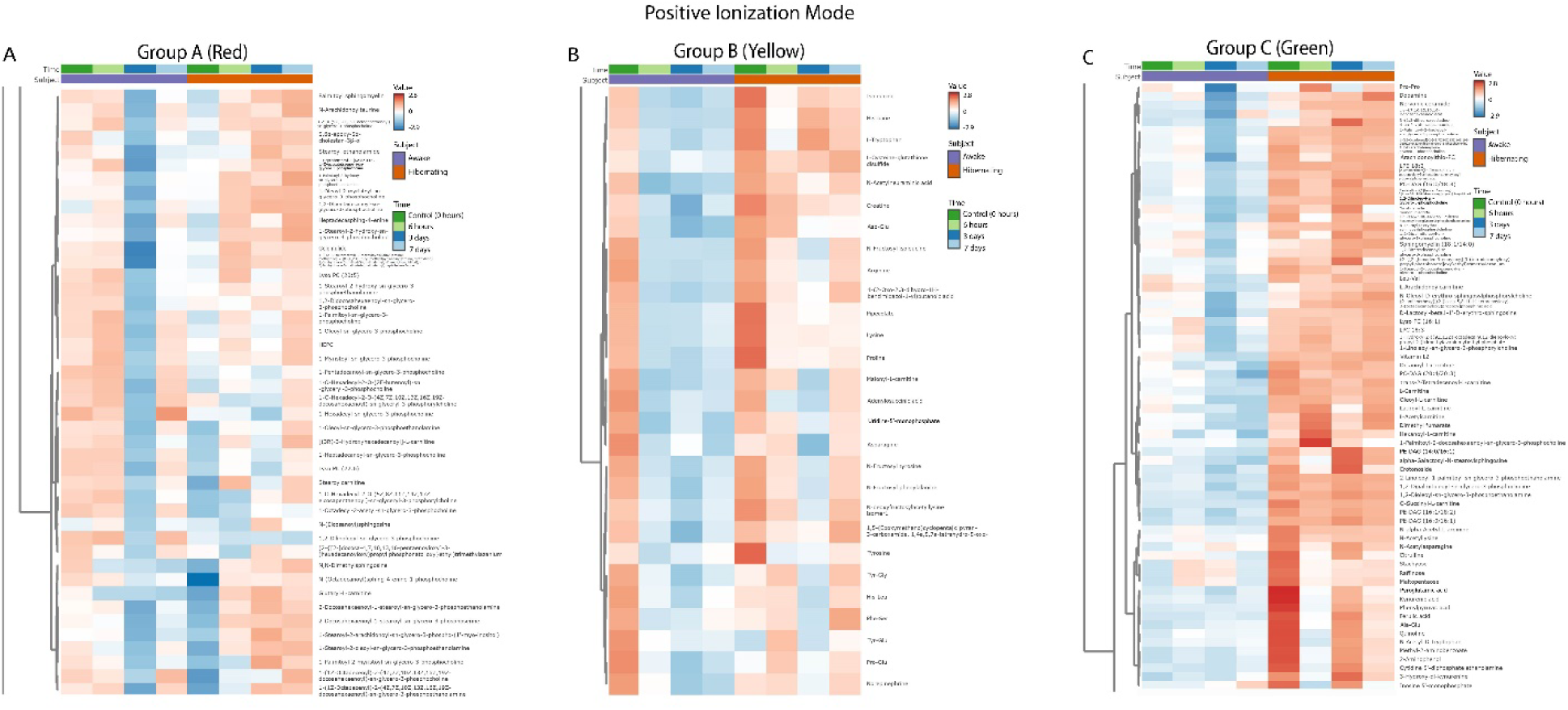
Heatmap clustering analysis of metabolites in the positive ionization mode for the three annotated groups (Group A-red, Group B-yellow, and Group C-green). Normalized feature intensity values ranged from -2.9 to 2.8. Subject 1 (S1): awake intact and injured TLGS; subject 2 (S2): hibernated intact and injured TLGS. Time 0 (T0): uninjured, intact control; time 1 (T1): 6 hours post-ONC; time 2 (T2): 3 days post-ONC; time 3 (T3): 7 days post-ONC.

Proline is particularly noteworthy due to its role in supporting RPE metabolism by fueling mitochondrial activity, contributing to the production of amino acids, and providing protection against oxidative stress [40]. Creatine, another metabolite elevated in hibernators, has demonstrated neuroprotective properties *in vitro* against both mitochondrial stress (e.g. sodium azide-induced inhibition of oxidative phosphorylation) and NMDA-mediated excitotoxicity. However, oral creatine supplementation in Sprague Dawley rats failed to confer neuroprotection in an ischemia-reperfusion model, likely due to downregulation of creatine transporters [41]. This suggests that although creatine influences energy metabolism, hibernation-induced neuroprotection likely involves a broader and more integrated metabolic program that cannot be fully replicated by creatine supplementation alone.

Elevated norepinephrine levels in Hibernating animals are also consistent with prior studies, as this neurotransmitter has been shown to protect neurons from microglia-induced activation and secondary cell death [42]. These findings suggest these metabolites may represent promising candidates for further investigation as protective and potentially therapeutic metabolites for retinal degenerative diseases, including glaucoma.

The second cluster, corresponding to Group B (green) identified in the negative ionization mode, highlights a set of metabolites that are elevated prior to injury and remain elevated following injury in Hibernating animals, but are consistently low in Awake squirrels (**Figure 5B**). This cluster includes nervonic ceramide, dopamine, and several types of metabolic acids such as pyroglutamic acid, kynurenic acid, and phenylpyruvic acid. The co-occurrence of kynurenic acid and tryptophan suggests activation of the kynurenine pathway, a branch of tryptophan catabolism known to modulate neuroinflammation and excitotoxicity. Kynurenic acid, in particular, is recognized for its neuroprotective effects and ability to protect against glutamate-induced excitotoxicity, a key contributor to RGC and neuronal injury [43, 44]. Also prominent in this cluster are various carnitines, which support mitochondrial function and buffer oxidative stress. Numerous DHA-containing phospholipids are included as well, many of which have been associated with protective roles in neuronal survival and membrane repair. Among the more unique metabolites elevated in hibernators is crotonoside, a compound extensively studied for its anti-tumor properties in acute myeloid leukemia [45] but only recently explored for its potential role in the central nervous system [46].

Additional metabolites of interest include dimethyl fumarate [47], a clinically approved treatment for multiple sclerosis, and ferulic acid [48] and quinoline [49], both of which have demonstrated neuroprotective effects in models of Alzheimer’s disease. Nicotinamide mononucleotide (NMN) is also elevated in hibernators even after injury, suggesting a critical role for NAD^+^ biosynthesis in RGC maintenance. NMN contributes to energy production via fatty acid oxidation [50] to stabilize ATP levels, reduce oxidative stress [51], and enhance DNA repair [52]. These functions are particularly relevant given that age-related NAD decline is associated with mitochondrial dysfunction and increased vulnerability to glaucomatous damage [53]. Another notable metabolite in this cluster is raffinose, a plant-derived oligosaccharide that appears elevated in hibernators and has been implicated in autophagy activation [54]. Though largely unexplored in the context of the central nervous system (CNS), its presence suggests a potential role in promoting cellular homeostasis and injury recovery.

Together, this broad repertoire of neuroprotective, anti-inflammatory, and stress-resilient metabolites underscores the systemic metabolic adaptations in hibernating squirrels. These adaptations likely contribute to the preservation of RGCs following optic nerve injury by supporting mitochondrial health, enhancing antioxidant defenses, modulating immune responses, and promoting DNA repair and autophagy.

The final cluster corresponding to Group C (red) was similarly defined by metabolites that declined at 3 days post-injury in Awake squirrels but increased in Hibernating squirrels at the same time point. This pattern highlights a set of potentially neuroprotective metabolites that are positively modulated by hibernation. Among these are several lipid mediators, including palmitoyl sphingomyelin, N-arachidonoyltaurine, and stearoyl ethanolamide—findings that align with observations from the negative ionization mode.

Also prominent in this cluster are multiple DHA-containing phospholipids associated with neuronal survival, antioxidant defense, and synaptic maintenance. These include 1,2-didocosahexaenoyl-sn-glycero-3-phosphocholine, 2-docosahexaenoyl-1-stearoyl-sn-glycero-3-phosphoethanolamine, and 2-docosahexaenoyl-1-stearoyl-sn-glycero-3-phosphoserine, with particularly robust elevations seen in the Hibernating TLGS retinas (**Figure 5C**).

Carnitine derivatives such as [(3R)-3-hydroxyhexadecanoyl]-L-carnitine—recognized for supporting mitochondrial function and offering neuroprotection in energy-compromised neurons—were also significantly elevated in Hibernating animals at 3 days post-injury. Additionally, the cluster contains lesser-characterized metabolites such as glycerophospholipids, which have been implicated in modulating neuroinflammation, and lysophosphatidylcholines containing EPA or DHA, which are known to support neuronal resilience. Together, the metabolites in Group C reinforce the observation that hibernation engages a metabolic program that elevates lipid-based neuroprotective signals, enhances mitochondrial support, and promotes a biochemical environment conducive to neuronal survival during peak injury stress.

### Pathway Analysis Reveals Hibernation Engages Amino Acid, Lipid, and Purine Pathways to Enhance Retinal Resilience

Pathway analysis for each subgroup was conducted in the positive and negative ionization modes using the complete set of significant metabolites in our data against the KEGG metabolic pathways (**Figure 6**). This analysis revealed a prominent enrichment in amino acid metabolism, as well as pathways related to the citric acid cycle, glycerophospholipid metabolism, and purine metabolism. During hibernation, metabolic activity is significantly depressed; however, these pathways provide alternative energy sources despite reduced oxygen and nutrient availability. Under metabolic stress, glycerophospholipids likely help maintain membrane fluidity and protect the central nervous system. Additionally, purine metabolism—including purine nucleosides such as adenosine, inosine, and guanosine—has been implicated in neuroprotection and is likely critical for regulating entry into and exit from torpor. Thus, hibernation triggers a cascade of adaptive responses that confer cytoprotection, which also serves to protect cells from trauma and injury, such as traumatic optic neuropathy.

**Figure 6.**
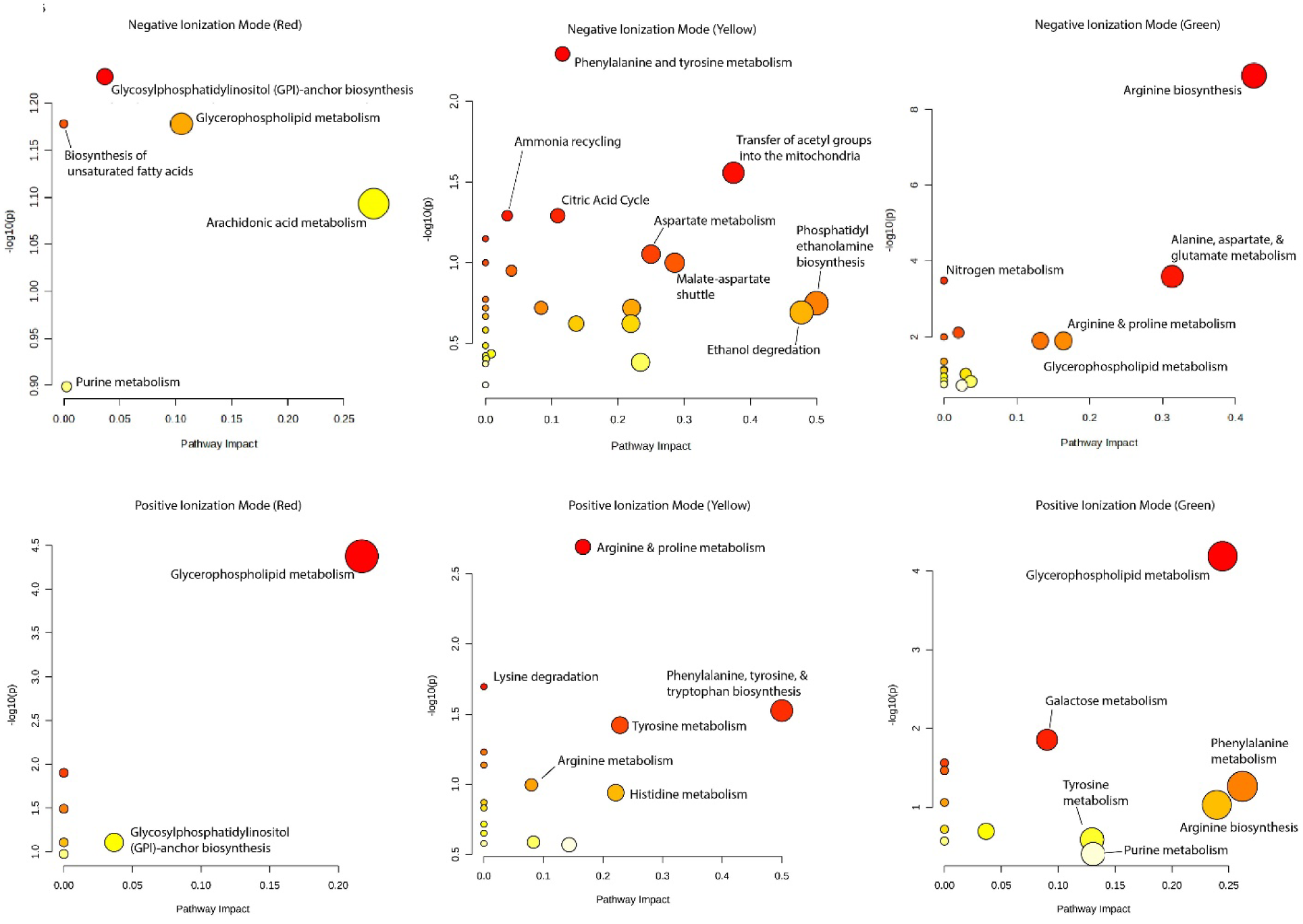
KEGG pathway enrichment analysis was performed using MetaboAnalyst 6.0 on all significantly changing metabolites (|log fold change| > 1.5) from the retina of Thirteen-lined Ground Squirrels (TLGS) with either intact optic nerves (ONs) or following optic nerve crush (ONC). The analysis identified significantly enriched metabolic pathways associated with injury-induced alterations in retinal metabolism.

### Validation of omics data implicating TLGS exosomes as neuroprotective

Hibernation is associated with metabolic signatures consistent with sustained increase in exosome biogenesis and release, suggesting a coordinated and adaptive mechanism for metabolic regulation and intercellular communication. The enrichment of exosome-related pathways during hibernation, as revealed by our omics data, highlights these vesicles as promising candidates for mediating neuroprotection through the transfer of metabolic substrates, signaling molecules, and antioxidant defense components. These findings support the hypothesis that exosome-based mechanisms, rather than isolated metabolic inhibition, underlie the robust neuroprotective phenotype observed in hibernating TLGS.

### Functional Characterization of TLGS-Derived Exosomes

We hypothesized that exosomes produced by TLGS cells modulate adaptive responses to cellular stress, including ON injury, thereby preventing neuronal death and axonal damage. To explore this possibility, exosomes were isolated from conditioned media of TLGS fibroblasts cultured at 37°C or 4°C for 48h. Isolated exosomes were ∼150nm in size, with concentrations of 2.8E10 particles/ml (**Figure 7, A-B**), and expressed canonical exosome markers, including CD81 (**Figure 7C**). Culture-derived exosomes provided a markedly higher yield than plasma-derived exosomes (**Figure 7C**). Isolation using the Exodus H-600 system minimized aggregates and debris, yielding high-purity preparations (**Figure 7D**).

**Figure 7.**
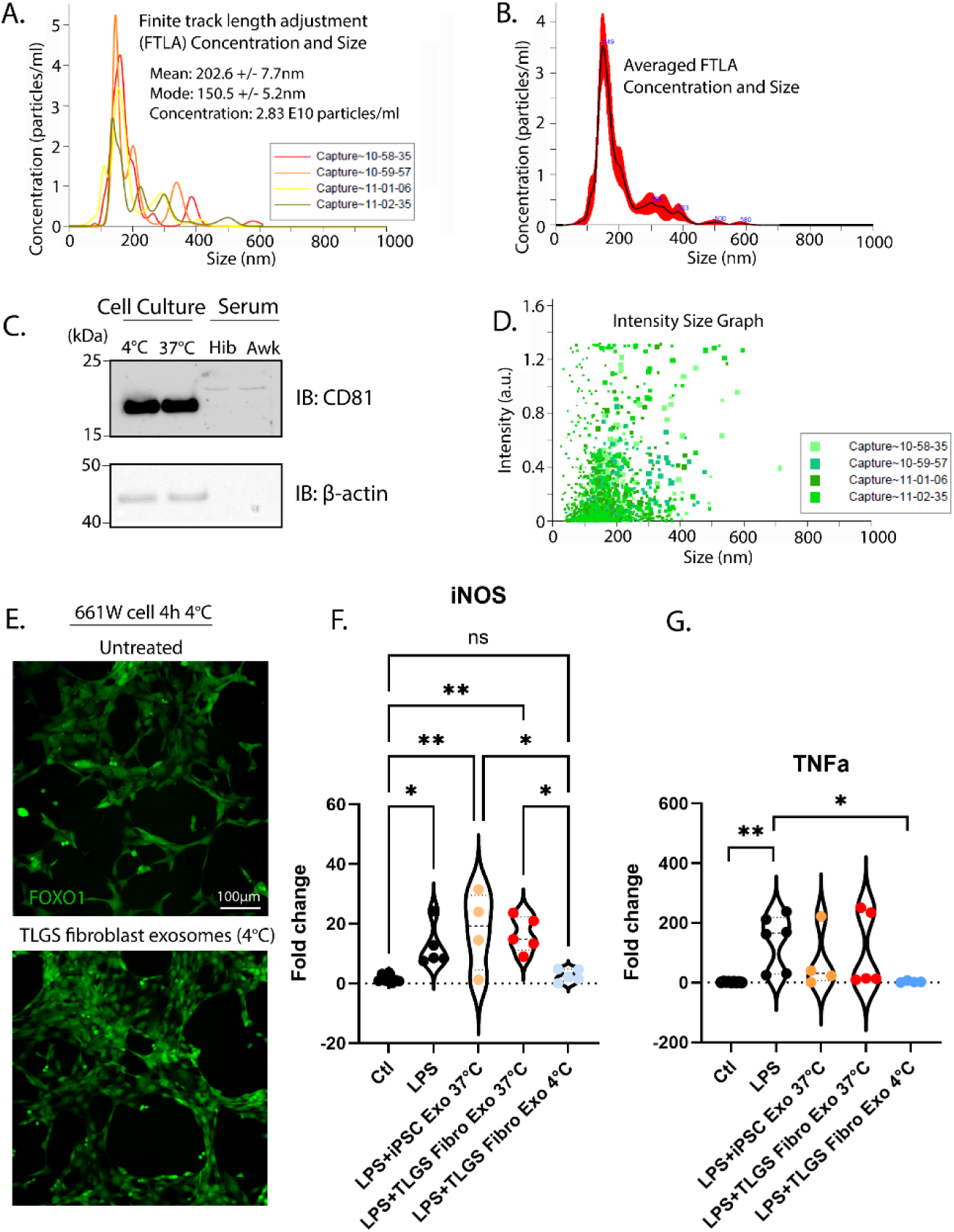
Neuroprotective and Anti-Inflammatory properties of TLGS exosomes. (A) Particle tracking analysis from four 30-s recordings demonstrates that iTLGS cells release 2.8 × 10¹ exosomes even at 4 °C. (B) Merged particle size distribution with a modal diameter of 148.7 nm. (C) Western blot confirming expression of the exosomal marker CD81. (D) Intensity vs. size plot, showing that larger particles scatter more light and dominate the profile; included as quality control to confirm minimal aggregate formation. E) Exogenous TLGS exosomes induced nuclear FOXO1 localization in neuronal cells from non-hibernating species, promoting survival under cold stress. F) Lipopolysaccharide (LPS)-induced activation of microglia leads to increased iNOS expression, which is reduced back to baseline following treatment with TLGS exosomes (4°C). G) TLGS exosomes (4°C) significantly suppress LPS-induced TNF-α expression, indicating anti-inflammatory potential. Significant changes are denoted by asterisks (*P ≤ 0.05, **P≤ 0.01).

In primary cultured TLGS fibroblasts, exosomes were essential for survival during temperature transitions, particularly during rewarming from 4°C to 37°C. A key component of this adaptive response was cold-induced FOXO1 nuclear translocation of FOXO1; however, non-hibernating mammalian cells appear to lose this capability with age [55]. Remarkably, treatment with exogenous TLGS-derived exosomes induced nuclear localization of FOXO1 in neuronal cells from non-hibernating species, promoting survival under cold stress conditions (**Figure 7E**). These findings suggest that exosome-mediated mechanisms of cold-induced neuroprotection can be transferred from hibernating to non-hibernating mammalian cells, conferring enhanced resistance to cellular stress.

### Neuroprotective and Immunomodulatory Effects of TLGS Exosomes

Functional assays demonstrated that exosomes from TLGS fibroblasts cultured at 37°C promoted survival of 661W photoreceptor cells under oxidative stress induced by H_2_O_2_ (**Figure 7F**). In BV2 microglial assays, pretreatment with cold-conditioned exosomes for 4 hours prior to 2-hour LPS exposure (0.2 μg/mL) significantly modulated inflammatory gene expression. RT-qPCR revealed marked reductions in iNOS and TNF-α mRNA compared to LPS-stimulated controls (**Figure 7G-H**). The immunomodulatory effects of cold-conditioned fibroblast exosomes exceeded those of exosomes from 37°C fibroblasts or TLGS iPSCs, indicating that both cell lineage and cold conditioning enhance the generation of pro-survival and anti-inflammatory factors.

### Omics Profiling of TLGS exosomes

Comparative LC–MS/MS proteomics (**Figure 8**) and whole transcriptomics analyses, including miRNAs (**Figure 9**) and mRNAs and LncRNAs (**Figure 10 A,B**) identified differentially enriched exosomal cargo (FC ≥ 1.5, adjusted p < 0.05) in primary cultured TLGS fibroblast cells exposed to 4°C compared with those maintained at 37°C. To prioritize functionally relevant and evolutionarily conserved cargo, candidates were ranked based on: 1. Evolutionary conservation across mouse and human orthologs, 2. Neurorelevance, including retinal/CNS expression and literature associations with glaucoma, neuroprotection, oxidative stress, or inflammation. Top conserved cargo included miRNAs miR-638 and miR-1269; mRNAs Naprt and Arhgef10l; and proteins HSPB1, PRDX6, and CLU. These components orchestrate multi-pathway neuroprotection in RGCs by: 1) maintaining redox balance (PRDX6, HSPB1, miR-638, Naprt), 2) stabilizing the cytoskeleton to prevent axonal collapse (Arhgef10l, HSPB1), and 3) regulating stress-response transcriptional programs (miR-1269, CLU).

**Figure 8.**
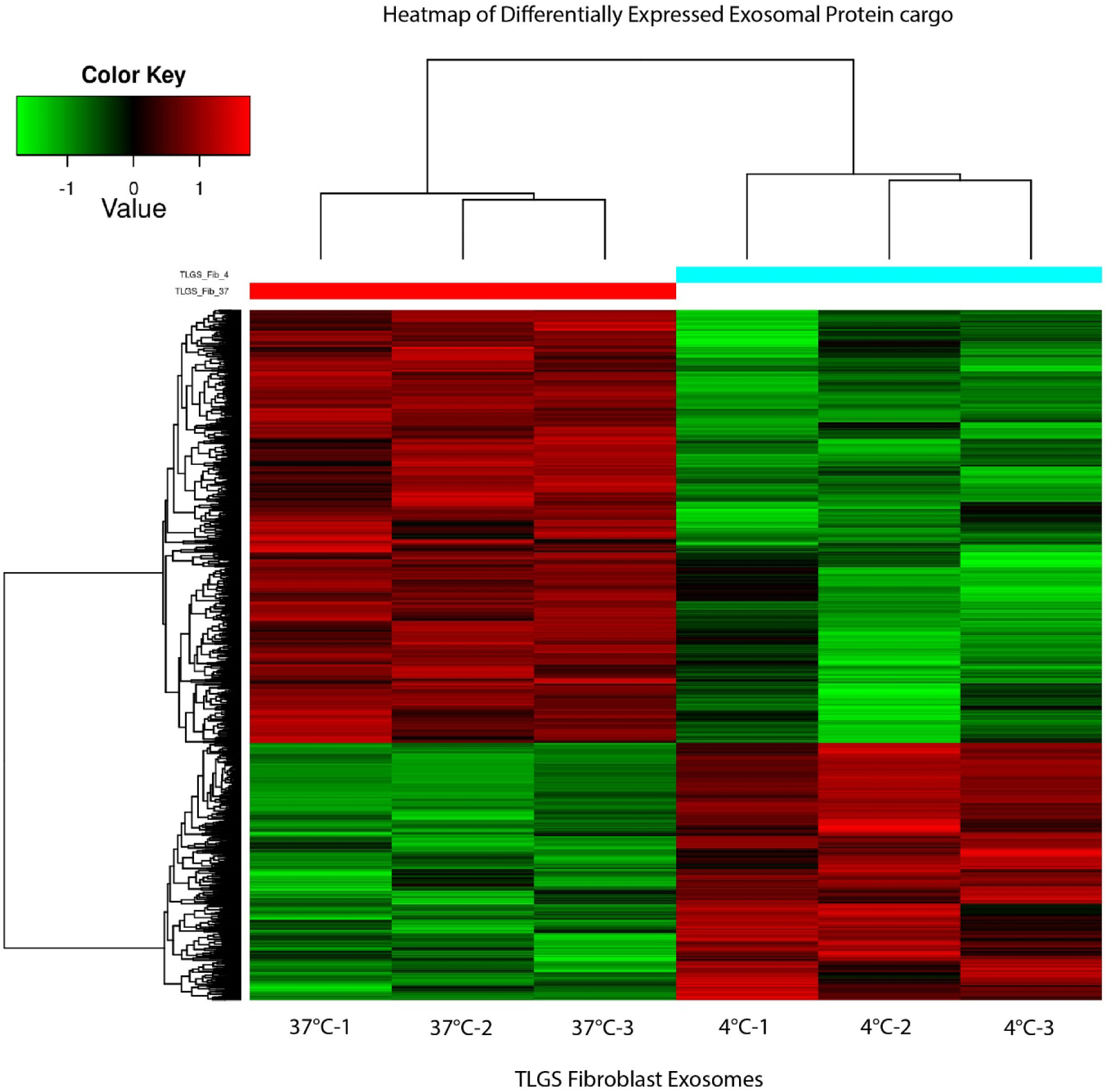
Heatmap of Differential Protein Clustering in exosomal cargo. The horizontal axis represents individual samples (37°C versus 4°C), and the vertical axis represents proteins showing significant differences between groups. Protein expression levels were standardized using the Z-score method and are displayed with a color gradient: red indicates significantly upregulated proteins, and green indicates significantly downregulated proteins. Protein names are provided in **Supplementary File S5**.

**Figure 9.**
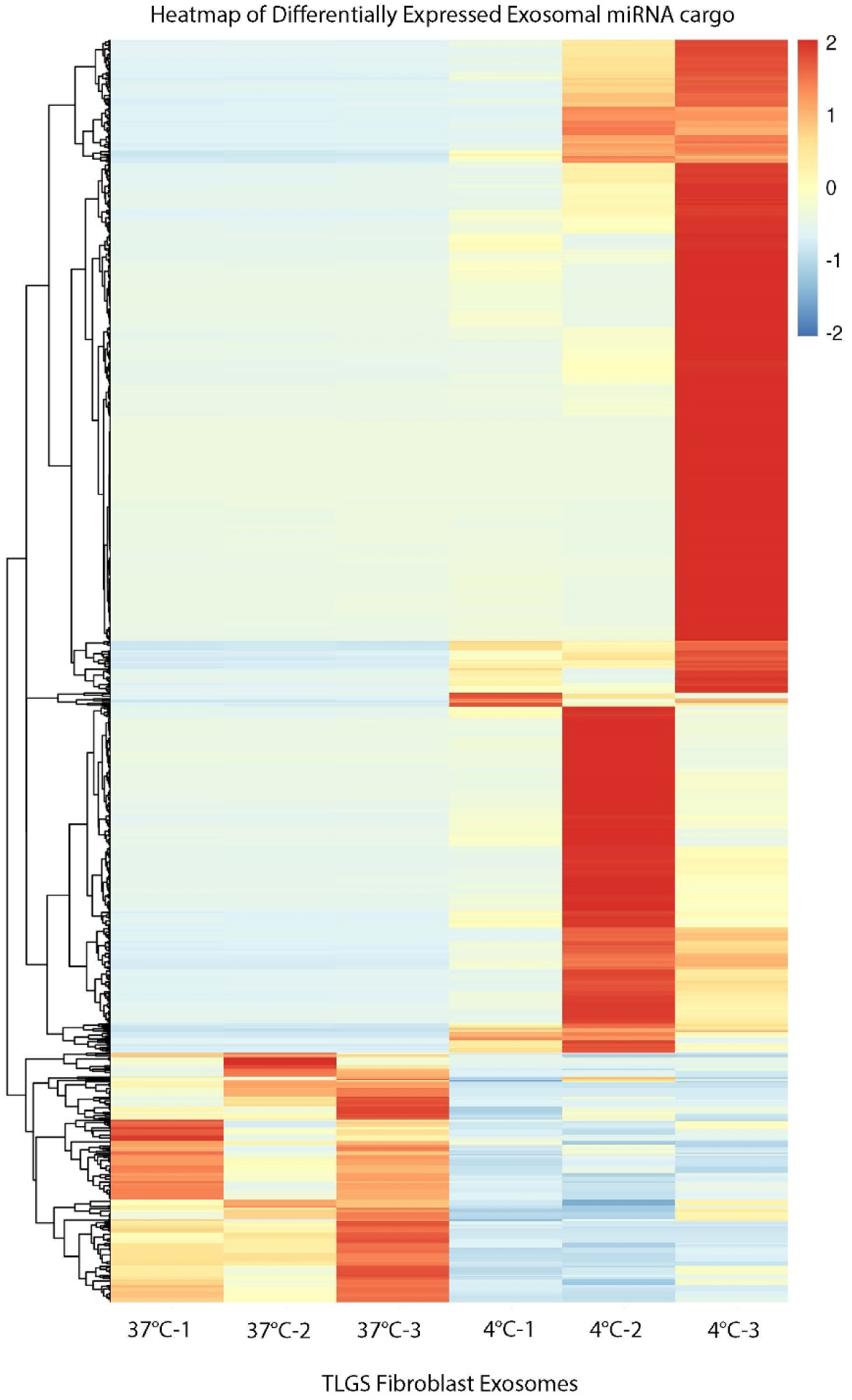
Heatmap of Differential miRNA expression. miRNA names are provided in **Supplementary File S6**.

**Figure 10.**
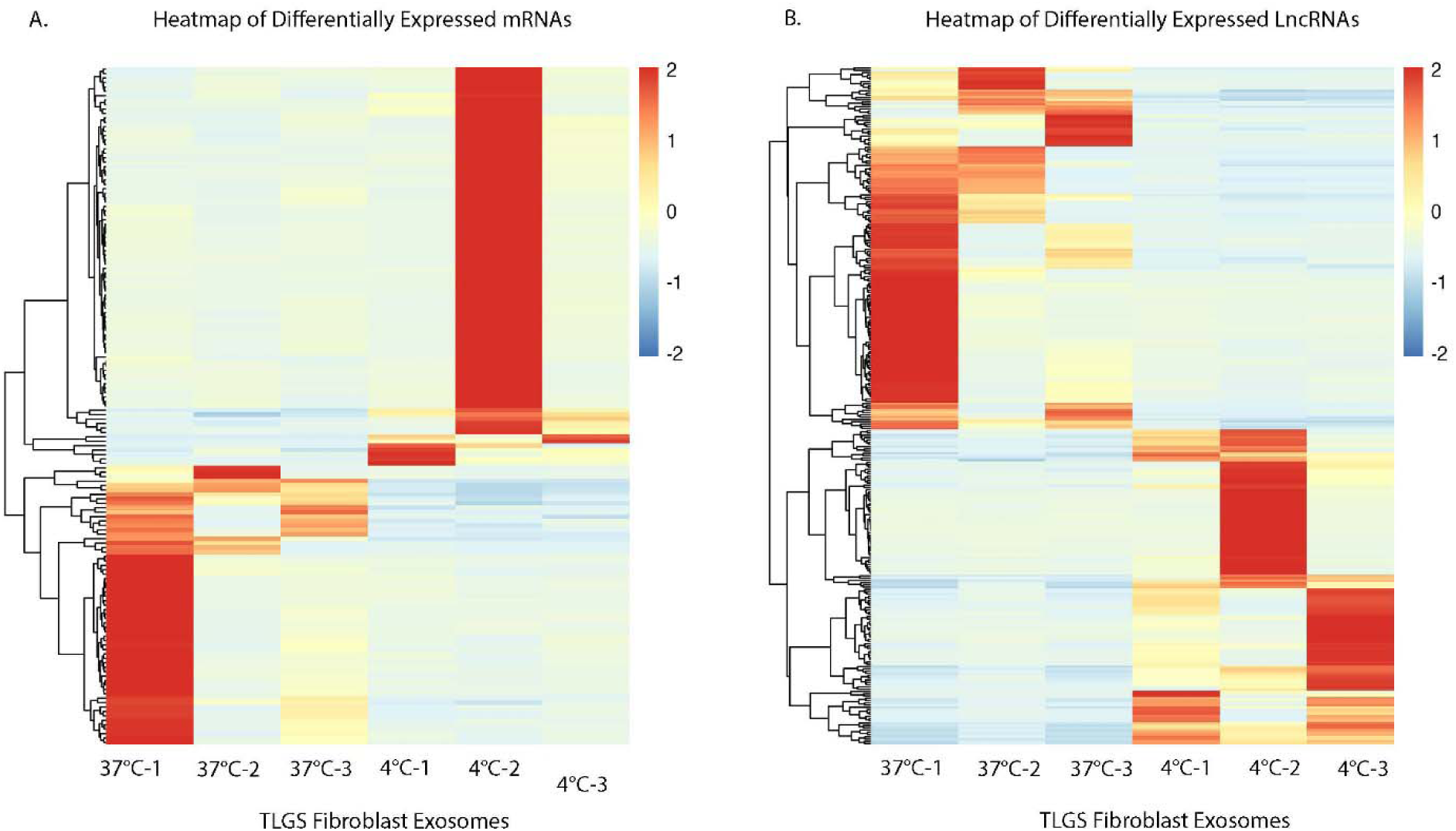
Heatmaps of Differential RNA Expression. (A) Differential mRNA expression and (B) differential lncRNA expression across samples. mRNA and lncRNA names are provided in **Supplementary File S7 and S8 respectively**.

Together, these findings validate that the neuroprotective phenotype observed in hibernating TLGS is not attributable to isolated metabolic inhibition, but rather to coordinated intercellular signaling mediated by exosomes. In contrast to pharmacological interventions that transiently modulate individual metabolic pathways while eliciting cytotoxic effects and disrupting the phagocytic clearance of dying cells [56], TLGS-derived exosomes confer broad neuroprotective benefits by promoting neuronal survival, suppressing inflammation, and restoring redox homeostasis under conditions relevant to optic nerve injury. Comparative omics profiling revealed that exosomes produced during cold adaptation are enriched in evolutionarily conserved miRNAs, mRNAs, and proteins that converge on key neuroprotective pathways, including redox regulation (PRDX6, HSPB1, miR-638, Naprt), cytoskeletal stability (Arhgef10l, HSPB1), and transcriptional control of adaptive stress responses (miR-1269, CLU). These integrated data support a model in which TLGS exosomes act as multifactorial mediators of neuroprotection—coordinating metabolic, structural, and immunoregulatory processes that collectively sustain RGC survival following injury.

## Discussion

Our metabolomics analysis revealed metabolites previously associated with optic nerve crush injury in non-hibernating species. Prior studies in mice have shown that ATP levels moderately increase within the first 10 hours post-ONC, likely due to enhanced mitochondrial oxidative phosphorylation [17]. Inhibition of ATP production—by blocking either glycolysis or oxidative phosphorylation in that model—increased RGC death. Notably however, supplying exogenous ATP had no effect on RGC survival. Based on these findings, the authors hypothesized that limiting oxidative phosphorylation could promote RGC survival in the ONC model. Indeed, treatment with meclizine, a drug that shifts energy metabolism toward glycolysis, resulted in improved RGC survival. These results suggest that increased mitochondrial respiration following ONC may contribute to ROS accumulation and secondary hypoxia, thereby exacerbating neuronal injury.

Consistent with these findings, our metabolomics data show a reduction in TCA cycle intermediates in Awake squirrels in response to ONC, suggesting enhanced mitochondrial respiration and increased TCA cycle activity in the injured retina. In contrast, hibernating animals exhibit an accumulation of TCA cycle intermediates, indicative of a metabolic shift toward alternative energy sources. Supporting this interpretation, we observed elevated levels of arachidonic acid—consistent with increased β-oxidation and lipid metabolism—as well as higher concentrations of carnitine derivatives, which facilitate fatty acid transport into mitochondria for oxidation. Additionally, increased glyceric acid, a product of glycerol oxidation during triglyceride breakdown, further supports a reliance on lipid-derived fuels. These metabolic changes align with previous reports describing adaptive energy strategies in hibernating species.[57–59]

A related metabolomic study examining lung preservation at 10°C identified loss of membrane lipid content as a potential limiting factor for extending transplant viability to 72 hours [60, 61]. Interestingly, in our hibernating samples, we observed increased levels of sphingolipids and ceramides, which may reflect a natural mechanism to replenish or stabilize membrane lipid content. While ceramide accumulation has been implicated in pathogenic processes under certain conditions (possibly influenced by whether the cells favor respiration or lipid metabolism), our data suggest a potentially protective role in the context of hibernation. This divergence may reflect differences in energy substrate utilization during torpor, where metabolic adaptations prioritize cellular survival under conditions of prolonged stress.

In hibernators, the maintenance of membrane fluidity appears to be a critical adaptive strategy, with downstream effects on lipid oxidation and metabolism. Membrane fluidity can influence the accessibility of lipids to oxidative enzymes, many of which are embedded in the lipid bilayer [62, 63]. Therefore, fluidity not only impacts the movement and interaction of these enzymes but also modulates their catalytic efficiency. Together, these findings point to a coordinated response in hibernators that preserves membrane integrity while supporting alternative metabolic pathways to promote cell survival under stress.

Our data identified upregulation of pathways involved in sphingolipid homeostasis, specifically ceramide biosynthesis, glutamine-glutamate cycle, and amino acid production. Metabolomic pathway analysis of this glaucoma model revealed increased levels of phenylalanine, tyrosine, tryptophan, arginine, and proline in hibernators. These amino acids paired with taurine as reported in our data may represent intrinsic mechanisms by which TLGS facilitate cell survival in response to glaucoma. Notably, tyrosine has previously been linked to glaucoma [64], while taurine and proline emerge as particularly promising candidates with potential therapeutic relevance in optic neuropathy.

In addition, the elevated levels of ceramide and sphingomyelins observed in Hibernating animals point to enhanced exosome-mediated signaling. These exosomes may carry neuroprotective, pro-survival, and anti-inflammatory cargo, highlighting hibernator-derived exosomes as a compelling avenue for future research and therapeutic development. Beyond their role in vesicular transport, ceramides and other sphingolipids are well-established modulators of immune signaling, particularly in glial cells. In microglia, ceramides can influence the balance between pro-inflammatory (M1-like) and anti-inflammatory (M2-like) activation states in a context-dependent manner that is closely linked to the cell’s metabolic profile. Therefore, annexin labeling in the retina following optic nerve crush [7] may serve as a diagnostic tool to assess membrane remodeling and apoptosis-associated signaling, potentially offering indirect insight into the retina’s underlying metabolic state.

Our metabolomic data reveal that TLGS exosomes carry cargo capable of transferring to non-hibernator cells and protecting them from oxidative stress. This highlights a cell-free, biologically inspired strategy to mimic hibernation’s protective effects, promoting cell survival through coordinated modulation of metabolism, inflammation, and neurodegeneration. TLGS exosomes or their contents could potentially be delivered via intravitreal or systemic administration, though optimization of dose, timing, and safety will be required. Our untargeted MS-based approach recognized metabolic changes that occur under both awake and hibernating conditions in TLGS after ONC. Statistically significant alterations were predominantly observed in metabolites directly associated with the tricarboxylic acid (TCA) cycle (**Figure 11**). Early steps of the TCA cycle exhibited significant changes that may contribute to downstream dysregulation of pathways such as ceramide biosynthesis, glutamine-glutamate metabolism, the purine nucleotide cycle, and nucleotide biosynthesis. These disruptions likely affect amino acid metabolism as well. However, it remains challenging to determine which pathways are initially dysregulated and thus drive subsequent metabolic disturbances, especially given the variation in injury timepoints. This represents a limitation of the present study and an important consideration for future investigations and also points to the potential usefulness of greater precision in these measurements, including the use of extraction controls and appropriate normalization procedures [65]. Another limitation lies in the relatively sparse annotation and identification of metabolites based on available codes, which constrains pathway enrichment analyses. Consequently, potentially important but poorly characterized metabolites may have been excluded from this analysis.

**Figure 11.**
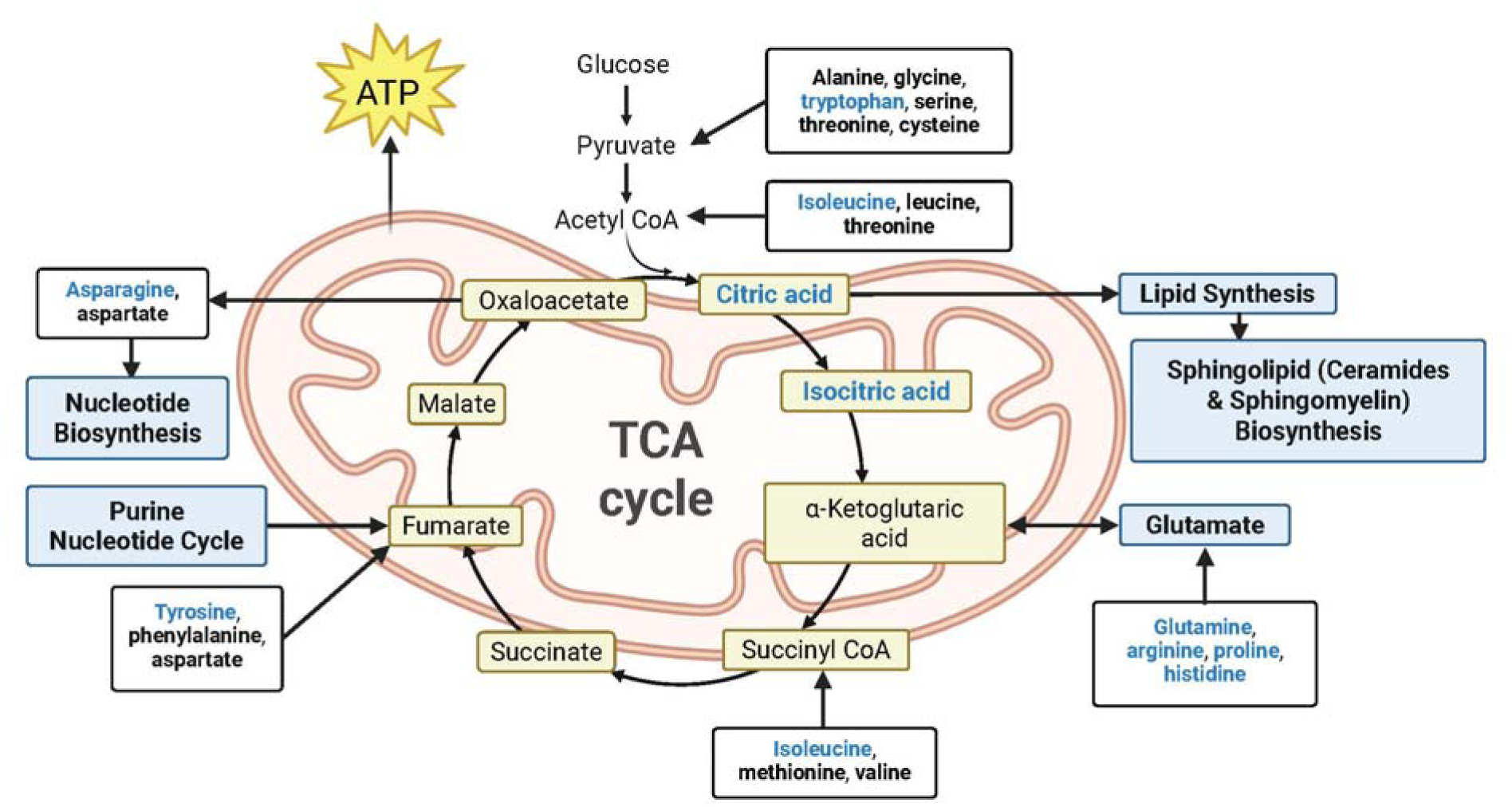
Schematic illustrating injury-induced metabolic changes in the retinas of awake Thirteen-lined Ground Squirrels (TLGS) and neuroprotective metabolic adaptations associated with hibernation. Metabolites shown in bold black were significantly dysregulated in Awake TLGS following optic nerve crush (ONC). Metabolites shown in bold blue were significantly decreased in Awake TLGS post-ONC but increased in Hibernating TLGS following ONC, suggesting potential hibernation-associated metabolic adaptations that support neuroprotection.

Anatomically, the TLGS retina more closely approximates the primate or human retina than other rodent models (mice and rats), based on RGC density and the organization of astrocytes along RGC axons. We acknowledge, however, that the TLGS model has limitations relative to human retinal physiology and disease, as even in the awake state these animals tend to be more resilient to cellular stressors, however, the hibernating state confers significantly more protection [66–68]. Additionally, while ONC provides a reproducible model of acute axonal injury, it does not fully recapitulate the pathophysiology of clinical conditions such as traumatic optic neuropathy or glaucoma, which involve chronic, multifactorial neurodegeneration. Therefore, characterization of closed-head impact model of traumatic optic neuropathy [69] and glaucoma in this species is warranted to determine their relevance to the human condition.

Despite these limitations, our study provides a comprehensive and nuanced metabolic profile of optic neuropathy in TLGS, highlighting key pathways that may mediate resilience to neural injury. The identification of hibernation-associated shifts—particularly in mitochondrial respiration, amino acid metabolism, and sphingolipid signaling—suggests that metabolic reprogramming is a core feature of neuroprotection in this model. These adaptive changes not only preserve membrane integrity and energy homeostasis but may also facilitate reparative signaling through exosome-mediated communication. Most notably, the convergence of altered TCA cycle activity, ceramide biosynthesis, and elevated neuroactive amino acids reveals a coordinated metabolic strategy that prioritizes cellular survival over rapid energy production. This is in stark contrast to the maladaptive responses observed in non-hibernating animals, where oxidative phosphorylation predominates and likely contributes to ROS-mediated damage. Taken together, our findings suggest that the metabolic state of the retina at the time of injury plays a critical role in determining neuronal fate. By uncovering the distinct metabolic signature of hibernation in the context of neuronal injury, we lay the groundwork for therapeutic approaches that emulate or induce these protective states. Approaches like hibernator-derived exosomes, ceramide-targeted interventions, and metabolic modulators may represent promising avenues to mitigate neurodegeneration and promote recovery following optic nerve trauma. Thus, hibernation may offer a powerful blueprint for reprogramming cellular metabolism to resist neurodegeneration—unlocking transformative strategies for treating optic neuropathies and other CNS injuries.

## Methods

### Study design

Retinas were collected from awake and hibernating thirteen-lined ground squirrels under the following conditions: for awake animals: 4 naïve, 12 subjected to total optic nerve crush injury (4 ONC 6h, 4 ONC 3d, 4 ONC 7d). For hibernating animals: 4 naïve, 12 subjected to total optic nerve crush injury (4 ONC 6h, 4 ONC 3d, 4 ONC 7d).

### Animals

All animal procedures were conducted in accordance with U.S. federal laws and Department of Agriculture regulations governing the humane use of animals in research and in adherence to the ARVO Statement for the Use of Animals in Ophthalmic and Vision Research (see https://www.arvo.org/About/policies/arvo-statement-for-the-use-of-animals-in-ophthalmic-and-vision-research/). Experimental protocols were reviewed and approved by the National Eye Institute Animal Care and Use Committee (NEI ASP-595). Adult male and female TLGS (*Ictidomys tridecemlineatus*), aged 6–12 months, were obtained from Dr. Dana Merriman (University of Wisconsin Oshkosh) and housed in the NIH animal facility under seasonally adjusted light–dark cycles that replicated natural photoperiods in Wisconsin. The optic nerve crush procedure is described in detail in the Supplementary Methods.

### Untargeted Metabolomics Sample Extraction

Samples were prepared first by homogenizing via bead-beating (Omni Hard Tissue Homogenizing Mix 2.8 mm ceramic (2 mL tubes), SKU 19-628; 50 mg biological material:1mL water; Omni Ruptor Bead Elite, 2.6 m/s, 30 seconds, 1 cycle); protein precipitation was then performed by adding ice-cold methanol (Fisher Chemical, Optima™ LC-MS grade) to homogenate (4:1 v/v). Samples were vortexed and placed in a –20°C freezer for 30 min to facilitate precipitation; samples were then centrifuged and 900uL of supernatant was transferred to a new 1.5 mL microcentrifuge tube. Samples were dried (Genevac EZ-2 Plus, 40°C, 2hr.) and resuspended via the addition of 200 microliters of water-acetonitrile 98%:2% v/v. An aliquot of each sample was transferred to a 2 mL autosampler vial with microvolume insert (Agilent); an aliquot from each extract was additionally transferred into a new 1.5 mL microcentrifuge tube to create a pooled quality control sample. Note: no extraction controls were used to assess recovery each time metabolites were recovered.

### Analytical Measurement

Samples were analyzed using an ultra-high performance liquid chromatograph (UHPLC, Vanquish, Thermo Scientific) coupled to a high-resolution mass spectrometer (Orbitrap Fusion Tribrid, Thermo Scientific) via heated electrospray ionization (NG Ion Max, Thermo Scientific) in positive and negative modes. UHPLC was used to separate chemicals, prior to mass spectrometric analysis, based on physical and chemical characteristics such as polarity. MS and MS/MS data were acquired. LC-MS data were collected from individual samples (n = 1 injection), system blanks (injection of solvent used to resolubilize samples), and a pooled quality control. The pooled quality control (QC) was injected multiple times at different volumes and used in data processing. MS/MS data, used to annotate features, were collected using the AcquireX (Thermo Scientific) deep scan methodology in which pooled QC was injected multiple times (n = 7). Chromatographic separation was carried out on a 2.1 x 100 mm, 100Å, 2.6 μm, F5 analytical column (Phenomenex) with corresponding guard cartridge. Gradient elution was performed using water with 0.1% acetic acid v/v and acetonitrile with 0.1% acetic acid v/v.

### Untargeted Metabolomics Data Processing

Compound Discoverer 3.3.0.550 (Thermo Scientific) was used to process data files which resulted in a tabular output of descriptors of each feature (e.g. m/z, retention time), annotation information (e.g. MS/MS database match), and peak area. We processed the output from Compound Discoverer using in-house R scripts via Jupyter Notebooks. The major components of the processing included formatting of the data outputs, comparison of m/z and retention time of annotation features versus an in-house generated list based on authentic chemical standards, assessment of signal response in pooled QC samples, assessment of signal variance in pooled QC samples versus samples (i.e. dispersion ratio), and multi- and univariate statistics. Features were annotated via spectral database matching to in-house libraries, NIST libraries, GNPS libraries, and m/zCloud libraries.

### MetaboAnalyst data analysis

Annotated features with assigned compound names were reviewed, and duplicate entries were removed. When duplicates were encountered, the feature with higher peak intensity and broader distribution across samples was retained. Compound names were standardized using the Metabolite ID Conversion tool available on MetaboAnalyst 6.0.[70] (https://new.metaboanalyst.ca/MetaboAnalyst/upload/ConvertView.xhtml). For any compounds that were not recognized by name or database identifiers, alternative names were queried using PubChem (https://pubchem.ncbi.nlm.nih.gov). Remaining unmatched compounds were subsequently searched using the LipidSig 2.0 ID conversion tool [71, 72] (https://lipidsig.bioinfomics.org/IDconversion/) to maximize annotation coverage. For visualization purposes, peak intensities from biological replicates within each experimental condition were averaged to simplify the display of data in principal component analysis (PCA) plots and heatmaps.

### Generation of Primary TLGS Fibroblasts

Primary fibroblast cultures were established from ear tissue obtained from deceased, euthanized thirteen-lined ground squirrels (TLGS). Hair was trimmed, and the tissue was sterilized by immersion in 70% ethanol for 15 minutes, followed by air-drying in a sterile plastic dish for 5 minutes. Using sterile forceps and a disposable scalpel, the tissue was minced into ∼1–2 mm² fragments and transferred into cryovials containing 0.25% trypsin. The vials were placed within 50 mL conical tubes and incubated in a rotating 37°C incubator for 40–45 minutes. The resulting cell suspension was filtered through a 100 µm cell strainer (Cole-Parmer, Catalog # UX-06336-71), and the tissue fragments were mechanically dissociated using the plunger of a sterile 3–5 mL syringe. Trypsin digestion was quenched by adding fibroblast culture medium consisting of DMEM supplemented with 5% fetal bovine serum (FBS), 1% penicillin–streptomycin, 1% sodium pyruvate, and 1% L-glutamine. The suspension was collected into a 15 mL tube and centrifuged at 800 rpm for 5 minutes. The resulting cell pellet was resuspended in fresh fibroblast medium and plated into a T-25 flask. Cells were allowed to adhere overnight, after which the medium was replaced. Cultures typically reached confluence within 3–5 days and were subsequently passaged for expansion.

### Exosome Isolation

TLGS cells were cultured in T-175 flasks until reaching approximately 80% confluence. The cells were then washed with 1x PBS and switched to 25ml of CO_2_-independent medium. Cultures were either incubated for an additional 48h at 37°C or transferred to a sterile fridge at 4°C. Conditioned media were collected in 50mL conical tubes and centrifuged at 2000 rpm for 20 min at 4°C (Eppendorf Centrifuge 5810R) to remove cell debris. The supernatant was transferred to new tubes and centrifuged again at 12,000 rpm for 30 min at 4°C using a Sorvall Lynx 6000 centrifuge. The resulting supernatant was carefully passed through a 0.2µm filter and stored in 50mL tubes at 4°C for short term storage or -80°C for long term storage. Fifty milliliters of conditioned medium were processed for exosome isolation using the H-600 Exosome Isolation System equipped with EID-MA03 cartridges (Exodus Bio). Following isolation, each cartridge was rinsed by pipetting 120–150 µL of sterile PBS up and down to elute the exosomes. The resulting eluate, containing the isolated exosomes, was stored at −80 °C until further use.

### Exosome Characterization using the Nanoparticle Flow Analyzer

Exosome samples were diluted 1:100 in sterile PBS and analyzed using a NanoSight NS-300 system (Malvern Panalytical) according to the manufacturer’s instructions. The “SOP Standard Measurement” protocol was used, consisting of five video captures, each 30 s in duration. Between each acquisition, the sample was briefly infused at 100 µL/min for 2–3 s and then stopped to advance exosomes into the field of view. During the first acquisition, infusion was halted as soon as nanoparticles appeared in the camera, allowing the instrument to focus on the exosomes. After all recordings were completed, data were processed by adjusting the “Detection Threshold” to optimally identify true particles; for this study, a threshold setting between 5 and 7 was used. Results were saved following completion of data processing.

### Exosome In vitro Neuroprotection Assays - XTT assay

661W cells were seeded into 96-well plates, and the XTT assay was performed according to the manufacturer’s instructions (Biotium, Catalog #30007). Two experimental conditions were used:

1. Oxidative stress model: Cells were pretreated with exosomes for 2 h, followed by incubation with H O for 6 h. The working XTT solution was prepared, and the activation reagent added. The activated XTT solution was then applied to the cells, which were incubated at 37 °C. Absorbance measurements were taken hourly for up to 3 h to monitor cell viability.
2. Cold stress model: Cells were pretreated with exosomes for 2 h, followed by incubation in OptiMEM supplemented with 10 mM HEPES at 4 °C for 16 h. The activated XTT solution was then added, and the plates were rewarmed at 37 °C for 2 h before absorbance measurements were recorded using a plate reader.

### Exosome Cargo Profiling - Proteomics and Whole Transcriptomics (WTS)

Exosome samples derived from 4 °C and 37 °C culture conditions were prepared in triplicate and submitted to Creative Biolabs for analysis. For whole transcriptome sequencing (WTS), at least 100 µL of exosome suspension per sample and replicate, with a concentration ≥10¹ particles/mL, was provided. For proteomic analysis, a minimum of 200 µL per sample and replicate, at the same concentration, was submitted. To prevent degradation and minimize freeze–thaw cycles, exosome samples were aliquoted according to analytical requirements and clearly labeled with the sample volume. Samples were suspended in sterile PBS, stored at −80 °C, and shipped on dry ice to maintain sample integrity during transport.

### Protein Extraction and Concentration Measurement

Exosome samples were thawed at 37 °C with gentle agitation, followed immediately by the addition of 5× RIPA lysis buffer. Samples were mixed thoroughly and lysed on ice for 30 min, with intermittent mixing. Protein concentration was determined using the bicinchoninic acid (BCA) assay. Briefly, 20 µL of each lysate was added to the BCA working reagent and mixed thoroughly. Samples were incubated at 37 °C for 30 min, and absorbance was measured at 562 nm using a spectrophotometer. Protein concentrations were calculated from a standard curve generated with known protein standards.

### Mass Spectrometry Analysis – Capillary High Performance Liquid Chromatography

Each sample is separated using a nanoflow HPLC liquid system. Buffer A is 0.1% formic acid aqueous solution, and Buffer B is 0.08% formic acid acetonitrile aqueous solution (80% ACN). The chromatographic column is equilibrated with 100% Buffer A, samples are loaded onto the mass spectrometry pre-column by an automatic sampler, then separated by the analytical column.

### Mass Spectrometry Identification

Each sample is analyzed by mass spectrometry using an Orbitrap Astral mass spectrometer (Thermo Scientific) after separation by capillary high-performance liquid chromatography (HPLC).

### RNA Extraction

RNA was extracted from exosome samples following the manufacturer’s protocol for the RNeasy Mini Kit (Qiagen, Catalog#: 217004) with slight modifications. Add 700 µL RNA lysis solution to the exosome sample, mix thoroughly, and incubate at room temperature for 5 min. Add 140 µL chloroform and vortex vigorously for 15 s to mix. Incubate at room temperature for 3 min, then centrifuge at 12,000 × g for 15 min at 4 °C (ensure the centrifuge is pre-cooled). Carefully transfer the upper aqueous phase to a new microcentrifuge tube, avoiding the interphase. Add 1.5 volumes of absolute ethanol (typically 525 µL) and mix by gentle pipetting. Load up to 700 µL of the mixture (including any precipitate) onto an RNeasy spin column and centrifuge at 8,000 × g for 15 s at room temperature. Discard the flow-through and repeat with any remaining sample. Add 700 µL Buffer RWT to the column, centrifuge at 8,000 × g for 15 s, and discard the flow-through. Wash the column with 500 µL Buffer RPE, centrifuge at 8,000 × g for 15 s, and discard the flow-through. Repeat the wash with 500 µL Buffer RPE, centrifuge at 8,000 × g for 2 min, and discard the flow-through and collection tube. Carefully remove the column to ensure no residual ethanol remains. Transfer the spin column to a new 2 mL collection tube and centrifuge at 12,000 × g for 1 min to dry the membrane. Discard the flow-through. Place the spin column in a new 1.5 mL RNase-free microcentrifuge tube, add 30 µL RNase-free water directly to the membrane, and centrifuge at 8,000 × g for 1 min to elute RNA. Store purified RNA immediately at −80 °C and record sample information.

### RNA Concentration Determination

RNA concentration was measured using a Quantus Fluorometer (Promega, E6150) according to the manufacturer’s instructions. Briefly, 1 µL of the extracted RNA was used for staining and quantification.

### Exosome RNA Qsep100 Analysis

RNA quality and size distribution were assessed using the Qsep100 system (BiOptic Inc.) with the NR1 cartridge. Based on the measured RNA concentration, an appropriate volume of RNA was diluted with the supplied diluent according to the manufacturer’s instructions, and the analysis was performed on the instrument.

### miRNA Library Construction

Adapters ligated sequentially to the 3’ and 5’ ends of miRNAs using the QIAseq miRNA Library Kit (Qiagen, Catalog #: 331505)

### cDNA Synthesis

cDNA was synthesized using the QIAseq miRNA NGS Kit (Qiagen) according to the manufacturer’s protocol.

#### Step 1 – Initial Reverse Transcription Setup

The following reaction system was prepared in each upstream PCR tube:

| Reagent | Volume (μL) |
| --- | --- |
| Upstream product | 40 |
| QIAseq miRNA NGS RT Initiator (RT) | 2 |
| **Total** | **42** |

The mixture was gently mixed and subjected to the following thermocycling conditions:

| Temperature | Time | Cycles |
| --- | --- | --- |
| 75 °C | 2 min | 1 |
| 70 °C | 2 min |  |
| 65 °C | 2 min |  |
| 60 °C | 2 min |  |
| 55 °C | 2 min |  |
| 37 °C | 5 min |  |
| 25 °C | 5 min |  |
| 4 °C | Hold |  |

#### Step 2 – Reverse Transcription Reaction

After the initial setup, the following reaction system was added directly to each PCR tube:

Step 2 – Reverse Transcription Reaction After the initial setup, the following reaction system was added directly to each PCR tube:
| Reagent | Volume (µL) |
| --- | --- |
| Upstream product | 42 |
| QIAseq miRNA NGS RT Primer | 2 |
| Nuclease-free Water | 2 |
| QIAseq miRNA NGS RT Buffer | 12 |
| QIAseq miRNA NGS RI | 1 |
| QIAseq miRNA NGS RT Enzyme | 1 |
| **Total** | **60** |

The mixture was gently mixed and subjected to the following reverse transcription program:

| Temperature | Time | Cycles |
| --- | --- | --- |
| 50 °C | 60 min | 1 |
| 70 °C | 15 min |  |
| 4 °C | 5 min |  |

### cDNA Purification using QMN Beads

Following reverse transcription, 143 µL of QMN Beads was added to each reaction, vortexed for 3 s, briefly centrifuged, and incubated at room temperature for 5 min. Tubes were centrifuged briefly and placed on a magnetic rack until the beads were fully separated, after which the supernatant was removed. Beads were washed twice with 200 µL of freshly prepared 80% ethanol without disturbance, then air-dried for 10 min with the lid open. Beads were resuspended in 17 µL of nuclease-free water, vortexed to mix, and incubated at room temperature for 2 min. After brief centrifugation and magnetic separation, 15 µL of the supernatant containing purified cDNA was transferred to a new microcentrifuge tube and stored at −20 °C until further use.

### Library Amplification

Library amplification was performed using the QIAseq miRNA NGS Library Kit following the manufacturer’s instructions. The PCR reaction mixture was prepared as follows:

| Reagent | Volume (μL) |
| --- | --- |
| Purified cDNA | 15 |
| Nuclease-free water | 42 |
| QIAseq miRNA NGS Library Buffer | 16 |
| HotStarTaq DNA Polymerase | 3 |
| Forward Primer | 2 |
| Index Primer | 2 |
| **Total** | **80** |

Reactions were gently mixed and subjected to PCR amplification under the following cycling conditions:

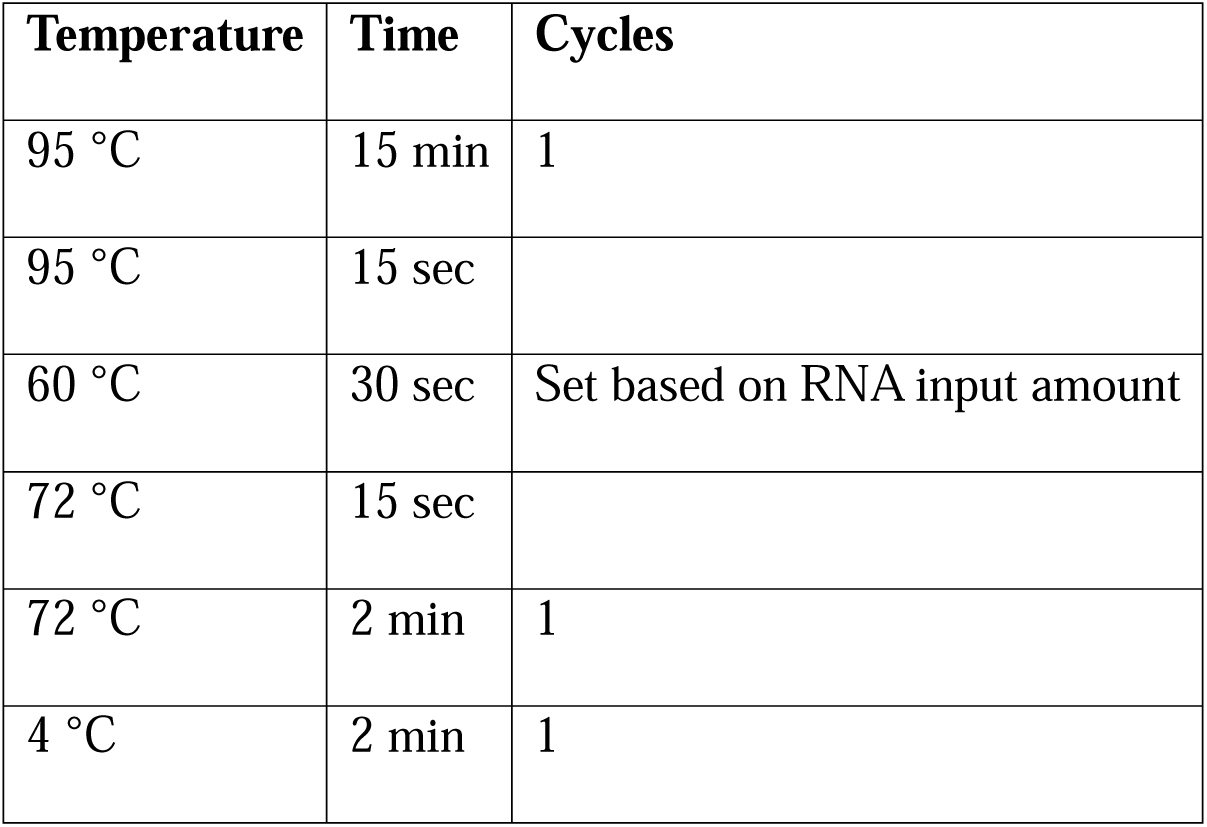
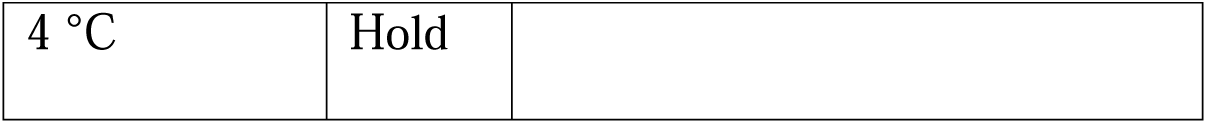

Following amplification, samples were either immediately subjected to library selection or stored at −20 °C until further use.

### Library Selection

Library amplification products were purified and size-selected using QMN Beads (BiOptic Inc.). Seventy-five microliters of the amplified library was mixed with 75 µL of QMN Beads, vortexed for 3 s, briefly centrifuged, and incubated at room temperature for 5 min. The tube was placed on a magnetic rack until the beads fully separated, and 145 µL of the supernatant was transferred to a new tube. Next, 130 µL of QMN Beads was added to the supernatant, vortexed for 3 s, centrifuged briefly, and incubated at room temperature for 5 min. After magnetic separation, the supernatant was carefully removed and discarded. Beads were washed twice with 200 µL of freshly prepared 80% ethanol, ensuring the pellet was not disturbed. Any residual liquid was removed, and the beads were air-dried for 10 min. The beads were resuspended in 17 µL of nuclease-free water, vortexed, and incubated at room temperature for 2 min. After magnetic separation, 15 µL of the supernatant containing the purified miRNA library was transferred to a new tube. An aliquot of the library was used for concentration and fragment size assessment, while the remaining library was stored at −20 °C until sequencing.

### RNA - Library Construction

#### Microscale Amplification

Between 0.5 and 4 µL of RNA sample was used for microscale amplification, with 13–15 PCR cycles (Thermo ProFlex PCR Machine) depending on input amount. Following magnetic bead purification, cDNA amplification products were quantified using the QuantiFluor dsDNA System (Promega, Catalog #: E2670).

#### Library Construction

One nanogram of cDNA was used for library construction with 13 PCR cycles. After bead purification, library concentration was measured using the QuantiFluor dsDNA System. An aliquot was diluted appropriately for fragment size analysis using the Qsep100 S2 cartridge (BiOptic Inc.).

#### Library Adapters and Sequencing

The structure and sequence of the library adapters are summarized in the table below. Libraries were sequenced using the Illumina NovaSeq 6000 platform in paired-end 150 bp (PE150) mode.

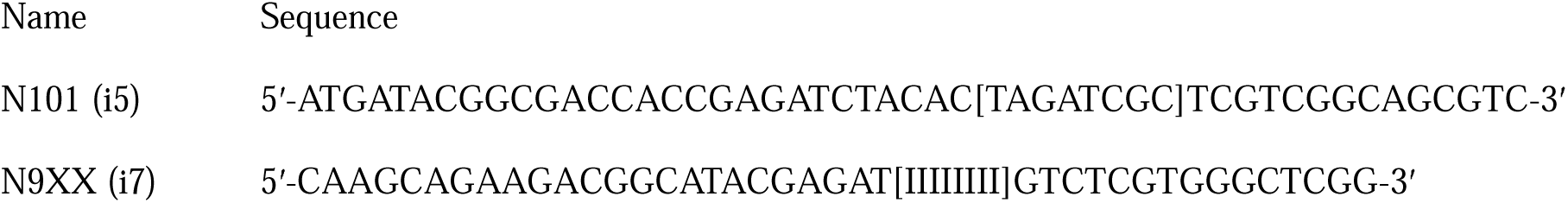

## Supporting information

Supplementary DataFile S1

Supplementary DataFile S2

Supplementary DataFile S3

Supplementary DataFile S4

Supplementary DataFile S5

Supplementary DataFile S6

Supplementary DataFile S7

Supplementary DataFile S8

## List of Abbreviations

ON: optic nerve
RGC: retinal ganglion cell
TLGS: thirteen-lined ground squirrel
CBT: core body temperature
ONC: optic nerve crush
h: hours
d: days
UHLPC: ultra-high performance liquid chromatograph
MS: mass spectrometry
MS/MS: Tandem Mass Spectrometry
LC-MS: Liquid Chromatography-Mass Spectrometry
QC: quality control
ROS: reactive oxygen species
iNOS: inducible nitric oxide synthase
TNF-α: tumor necrosis factor alpha
Iba1: ionized calcium-binding adaptor molecule 1
CD68: Cluster of Differentiation 68
GFAP: Glial Fibrillary Acidic Protein
DAPI: 4′,6-diamidino-2-phenylindole
PLS-DA: partial least squares discriminant analysis
HIF-1α: Hypoxia-Inducible Factor 1 alpha
Cer: ceramide
SM: sphingomyelin
AMD: age-related macular degeneration
mRNA: messenger ribonucleic acid
microRNA: micro ribonucleic acid
LPE: lysophosphatidylethanolamine
Nrf2: Nuclear Factor Erythroid 2-Related Factor 2
RPE: retinal pigment epithelium
NMDA: nictotinamide
NMN: Nicotinamide mononucleotide
ATP: adenosine triphosphate
DNA: deoxyribonucleic acid
CNS: central nervous system
EPA: eicosapentaenoic acid
DHA: docosahexaenoic acid
TCA: tricarboxylic acid

## Ethics approval and consent to participate

All animal experiments were approved by the Institutional Animal Care and Use Committee of the National Eye Institute (Animal Study Protocol: NEI-595).

## Consent for publication

Not applicable

## Availability of data and Materials

The datasets supporting the conclusions of this article are included within the article (and its additional files). The raw untargeted metabolomics data is available on MassIVE (ID#: MSV000099893).

## Authors’ contributions

KJM: Funding acquisition, Conceptualization, Methodology, Formal Analysis, Visualization, Investigation, Supervision, Writing - Original Draft, Writing-Reviewing and Editing. FMN: Funding acquisition, Conceptualization, Methodology, Formal Analysis, Visualization, Investigation, Writing-Reviewing and Editing. RM: Writing - Original Draft, Methodology, Formal Analysis, Visualization, Investigation, Writing-Reviewing and Editing. KO: Investigation, Formal Analysis, Visualization, Writing - Review & Editing. AJ: Investigation, Formal Analysis, Visualization, Writing - Review & Editing.

## Acknowledgements

The authors thank the NIH Division of Veterinary Resources for providing veterinary care and technical research support for the TLGS colony and to Dr. Dana Merriman (University of Wisconsin, Oshkosh) for providing the TLGS used in the study. The authors also thank Dr. John Ball for providing the body temperature trace from a hibernating thirteen-lined ground squirrel, which is presented in Figure 1B and a single interbout arousal is shown at higher temporal resolution in Figure 1C.

## Author information

Francisco M. Nadal-Nicolas, Rachel McNeel, Kiyoharu J. Miyagishima contributed equally to this work.

## Supplementary Material

**Supplementary Figure S1.**
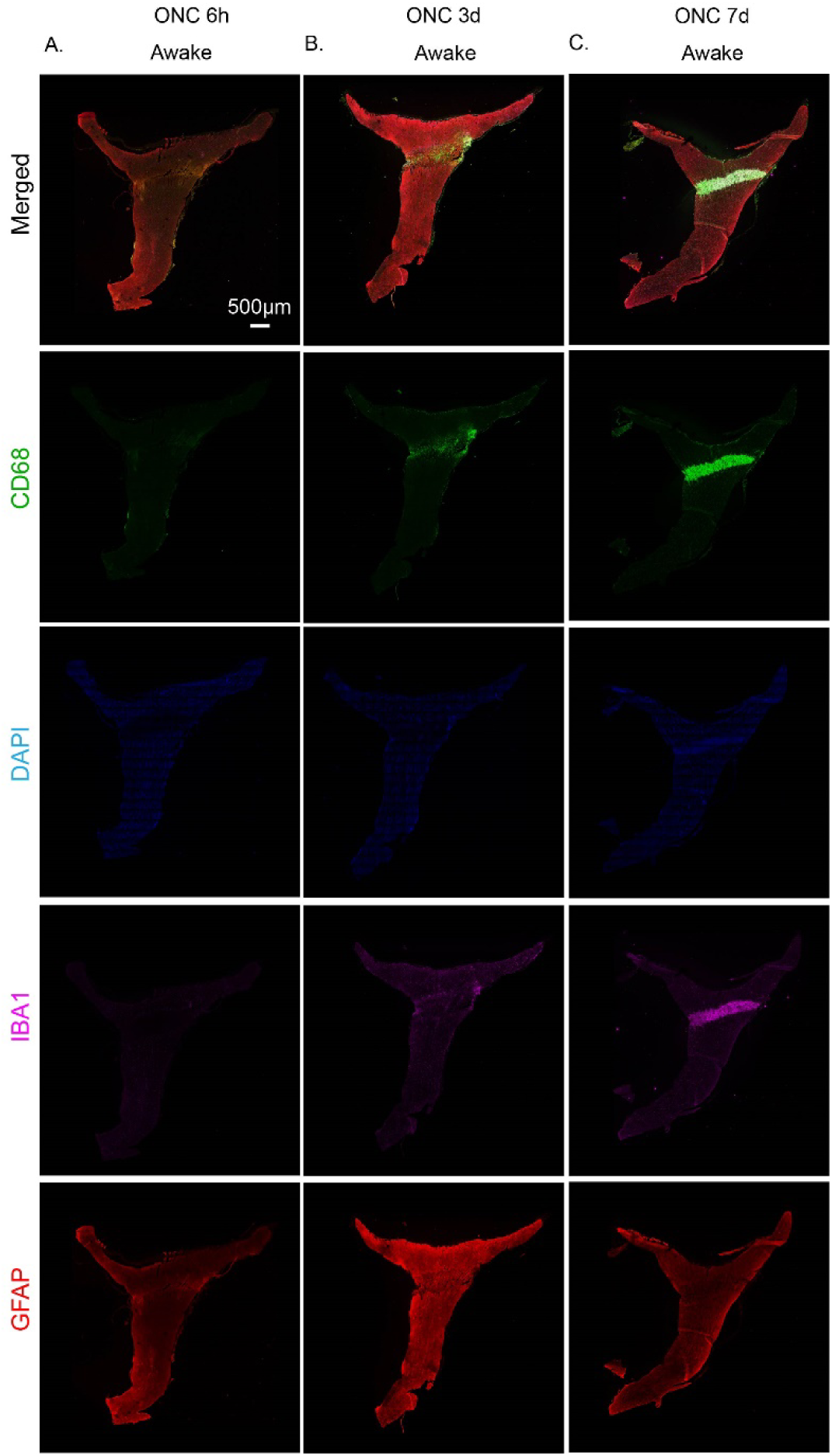
Confocal images of representative optic nerves from Awake TLGS following optic nerve crush, collected at 6 h (A), 3 d (B), 7 d (C) post-injury. Optic nerves were stained for CD68 (green), IBA1 (magenta), GFAP (red), and counterstained with DAPI (blue). Scale bar: 500 µm.

**Supplementary Figure S2.**
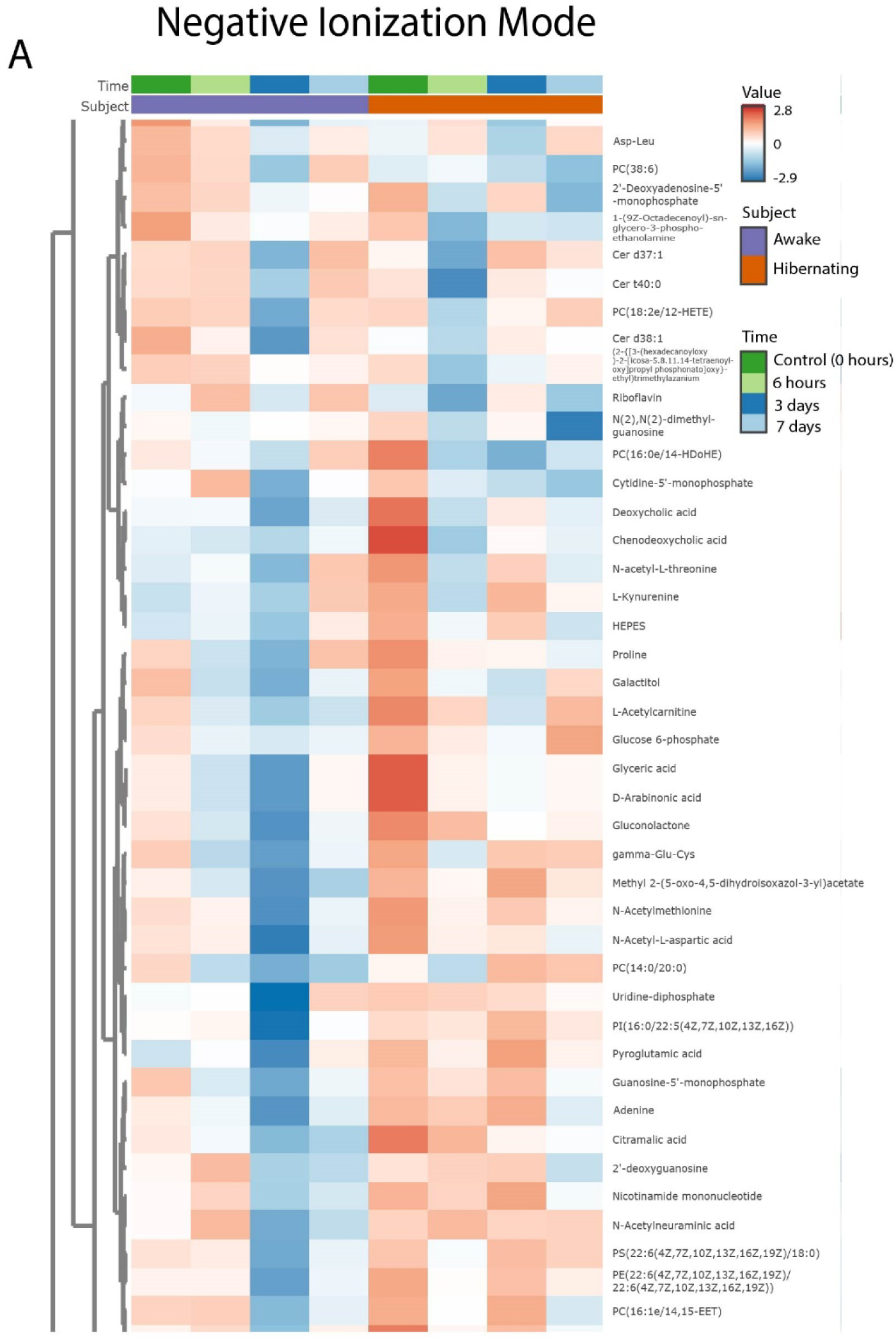
Heatmap of additional metabolites identified in negative ionization mode. Metabolites in this group exhibit variable changes following injury.

**Supplementary Figure S3.**
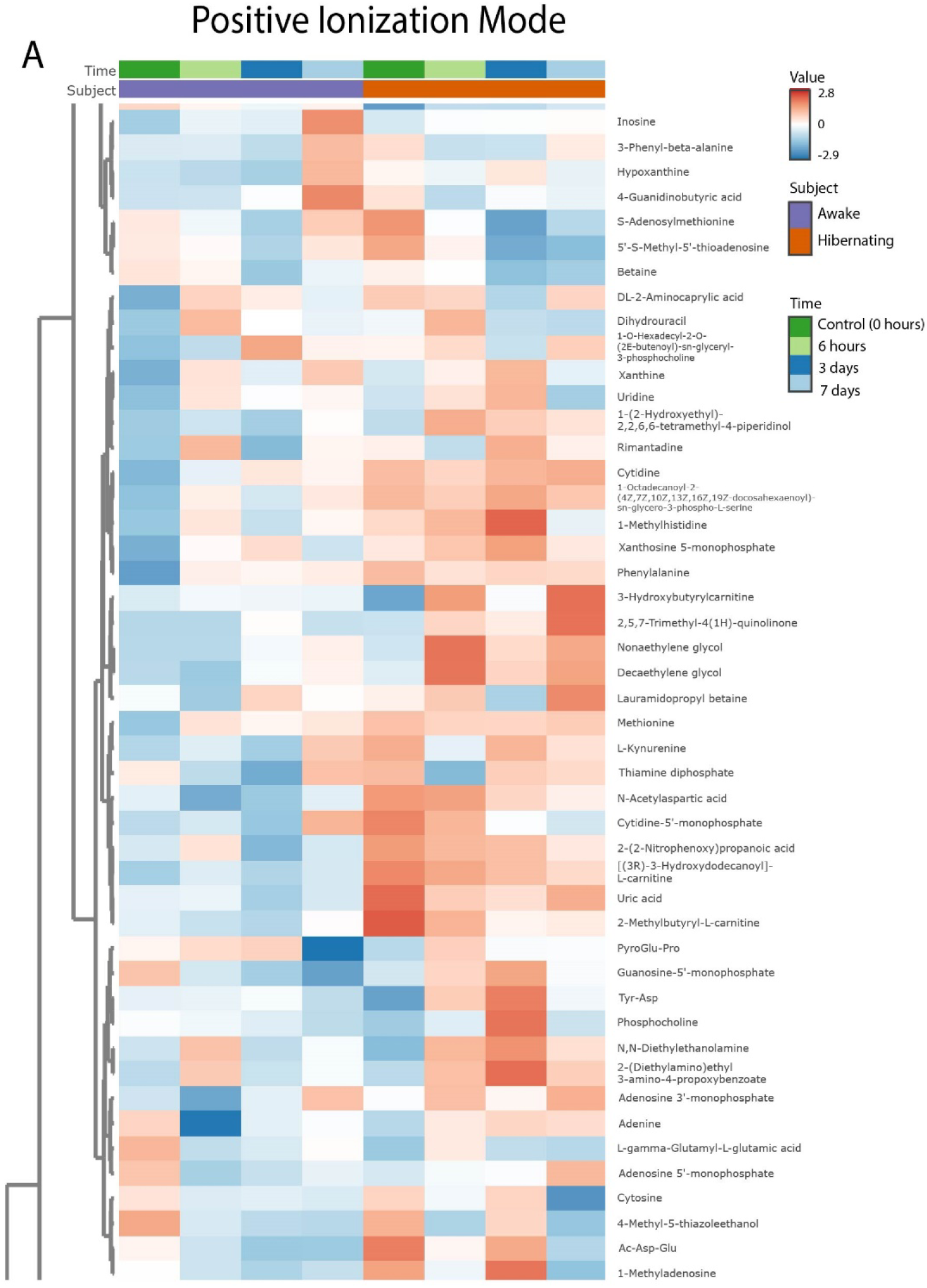
Heatmap of additional metabolites identified in positive ionization mode. Metabolites in this group exhibit variable changes following injury

## Supplementary Methods

### Optic Nerve Crush

Squirrels were anesthetized using isoflurane gas, with induction at 5% and maintenance at 4% via a low-profile nose cone. Anesthetic gas was scavenged using an attached activated charcoal filter. Sterile surgical instruments were autoclaved prior to use and subsequently disinfected between animals by immersion in 70% ethanol followed by bead sterilization at approximately 270°C for 20 seconds. To prevent infection, topical antibiotic (polymyxin B-neomycin-bacitracin) ophthalmic ointment was applied to the surgical eye. To prevent corneal dehydration during anesthesia, Systane® lubricating gel was applied to both eyes. Ketoprofen (5 mg/kg, subcutaneous) was administered prior to recovery to provide preemptive analgesia, which helps reduce the severity and duration of postoperative pain.

For hibernating animals, body temperature was maintained at approximately 4°C using an ice pack placed beneath the animal during the procedure. Animals were anesthetized with isoflurane (induction at 3%, and maintenance at 3%) to sustain the hibernation state and prevent arousal. Animals were kept under isoflurane anesthesia to maintain the hibernation state and prevent arousal. To minimize sensory stimulation, ambient lights were turned off, the room was isolated from external noise, and exposure to illumination from the surgical microscope was kept to a minimum. To maintain body temperature during transfer and to minimize light, animals were transferred to and from the hibernaculum in a foam shipping container equipped with ice packs. In the fall preceding the hibernation season, animals were implanted subcutaneously with temperature transmitters (TA-F40, Data Sciences International) or iButtons (AlphaMach), enabling wireless body temperature monitoring to confirm that animals remained in torpor following the surgical procedures [73].

For awake, normothermic animals, a disposable hand warmer wrapped in a paper towel was used to provide gentle warmth while preventing direct skin contact.

Under a dissection stereo microscope, a temporal canthotomy was performed on one eye, followed by a lateral conjunctival incision near the corneal margin. The retinal blood supply was preserved throughout the procedure. After gently separating the retractor bulbi muscles, the optic nerve was exposed intraorbitally using blunt dissection. A calibrated cross-action forceps was used to compress the optic nerve for 10 seconds, approximately 1–2 mm posterior to the globe.

After the injury was induced, the surgical site was closed with sutures. Animals were monitored continuously throughout recovery from anesthesia and returned to their home cages once fully ambulatory. Postoperative monitoring included assessment for signs of pain or distress. Any animal exhibiting signs of discomfort was evaluated in consultation with veterinary staff and treated or euthanized as necessary.

Experimental endpoints for the metabolomics study were at 6 hours, 3 days, or 7 days post-injury. For anatomical assessments, additional survival time points included 14 days post-injury.

### Immunohistochemistry: Retina

Eyes were fixed in 4% PFA for 1 hour followed by dissection in PBS. Using a binocular dissecting scope, the anterior part of the eye is removed by first making an incision with a razor blade and then inserting dissection scissors to remove the lens and cornea. The eyecup is then cut into a 4-petal flower shape by making radial incisions at 90-degree intervals. Using fine forceps, the retina was carefully lifted away from the underlying retinal pigment epithelium. Special care must be taken to gently cut the retina free from the optic nerve head as the optic nerve in squirrels is oriented horizontally.

After carefully removing the vitreous using fine forceps and a soft brush, retinas were permeabilized and blocked in 2 % normal goat serum (NGS) with Triton X-100 in PBS for 30 min, replacing the solution every 10 min [74]. Tissues were then incubated at 4 °C for 7 days with a primary antibody against antibrain-specific homeobox/POU domain protein 3A (Brn3a, red, 1:750, Cat#: C-20 [discontinued]; Santa Cruz Biotechnologies) or RNA Binding Protein, Multi-Splicing (RBPMS, red, 1:500, Cat#: GTX118619; Genetex). Following thorough PBS washes, retinas were incubated with the appropriate Cy3-conjugated donkey secondary antibody overnight at room temperature.

### Immunohistochemistry: Optic nerve

Optic nerves were fixed in 4% PFA for 1 hour at room temperature, followed by three washes in PBS. Fixed tissues were embedded in OCT compound and 30-um thick cryosections were collected using a cryostat (Leica, Nussloch, Germany). Cryosections were incubated with antibodies for CD68 (green, Biorad, Cat#: 019-19741, 1:100), IBA1 (magenta, Wako, Cat#: 019-19741, 1:100), and glial fibrillary acidic protein (GFAP, red, Aves Labs, Cat#: GFAP87987979, 1:500). Nuclei were counterstained with DAPI (blue, 1:2000). After staining, sections were mounted on glass slides using antifade mounting medium and imaged using a Zeiss LSM 780 confocal microscope.

### Gene expression analysis

RT-qPCR was used to quantify gene expression changes in BV2 microglial cells. BV2 microglial cells were treated with various reagents (e.g., LPS or treated with TLGS-derived exosomes) for 6 hours, after which cells were collected for RNA extraction and subsequent RT-PCR analysis to quantify gene expression changes. Total RNA was extracted using NuceloSpin RNA, Mini kit for RNA purification (Takara Bio, Cat: 740955.50) and reverse-transcribed to cDNA using (Takara PrimeScript 1^st^ strand cDNA synthesis kit (Takara Bio, Cat: 6110A) according to manufacturer’s instructions. cDNA was used for RT-qPCR in a reaction volume of 15µl containing, 2µL cDNA, 2µl of primer mixture, and 7.5 µl of Master mix (EmeraldAmp® GT PCR Master Mix, Takara Bio, Cat: RR310B) and 3.5 µl of water. RT-qPCR was performed using a CFX96 Real-Time PCR Detection System (Bio-Rad). Ribosomal protein S14 (RPS14) was used as an internal control. mRNA expression levels were normalized to control samples using the comparative Ct (2^−ΔΔCt^) method. A list of primers is shown below. At least 3 replicates of each experiment were performed.

Primers:

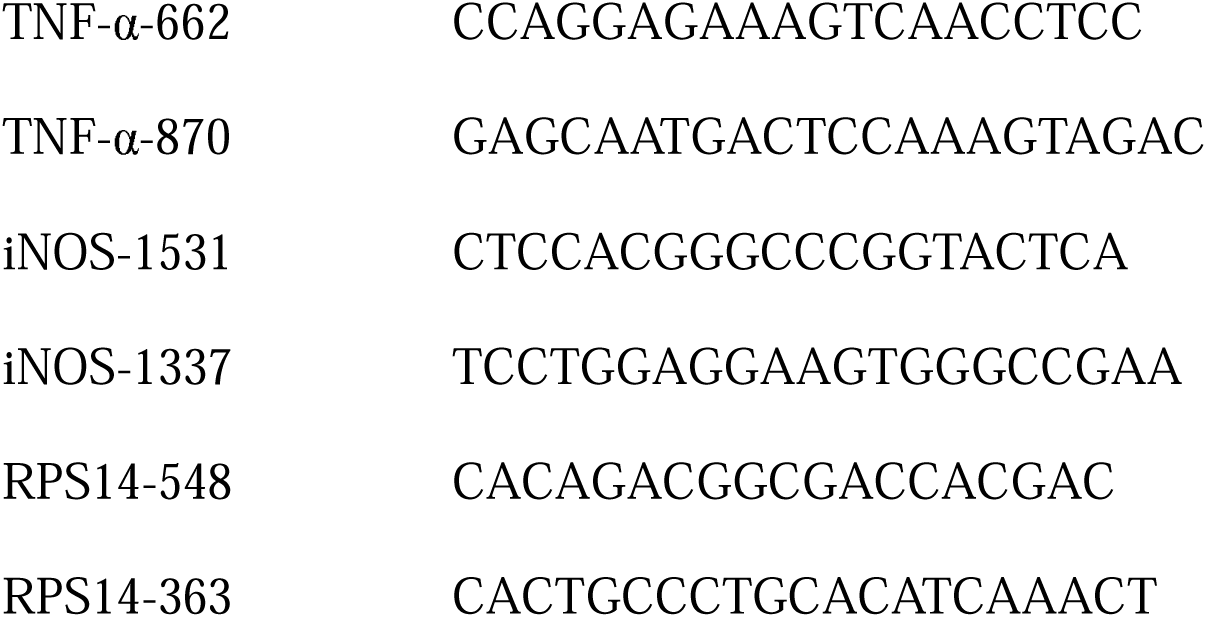

### Statistical analysis

Statistical analyses were performed using GraphPad Prism, Microsoft Excel, or custom MATLAB scripts. Pairwise comparisons were conducted using paired t-tests, while comparisons among three or more groups were performed using one-way analysis of variance (ANOVA).

